# Ectopic FVIII expression and misfolding in hepatocytes as a cause for hepatocellular carcinoma

**DOI:** 10.1101/2021.01.19.427220

**Authors:** Audrey Kapelanski-Lamoureux, Zhouji Chen, Zu-Hua Gao, Ruishu Deng, Anthoula Lazaris, Cynthia Lebeaupin, Lisa Giles, Jyoti Malhotra, Jing Yong, Chenhui Zou, Ype P. de Jong, Peter Metrakos, Roland W. Herzog, Randal J. Kaufman

## Abstract

Hemophilia A gene therapy targets hepatocytes to express B domain deleted-(BDD) clotting factor VIII (FVIII) to permit viral encapsidation. Since BDD is prone to misfolding in the endoplasmic reticulum (ER) and ER protein misfolding in hepatocytes followed by high fat diet (HFD) can cause hepatocellular carcinoma (HCC), we studied how FVIII misfolding impacts HCC development using hepatocyte DNA delivery to express three proteins from the same parental vector: 1) well-folded cytosolic dihydrofolate reductase (DHFR); 2) BDD-FVIII, which is prone to misfolding in the ER; and 3) N6-FVIII which folds more efficiently than BDD-FVIII. One week after DNA delivery, when FVIII expression was undetectable, mice were fed HFD for 65 weeks. Remarkably, all mice that received BDD-FVIII vector developed liver tumors, whereas only 58% of mice that received N6 and no mice that received DHFR vector developed liver tumors, suggesting that the degree of protein misfolding in the ER increases predisposition to HCC in the context of a HFD and in the absence of viral transduction. Our findings raise concerns of ectopic BDD-FVIII expression in hepatocytes in the clinic, which poses risks independent of viral vector integration. Limited expression per hepatocyte and/or use of proteins that avoid misfolding may enhance safety.

**Graphical abstract:** 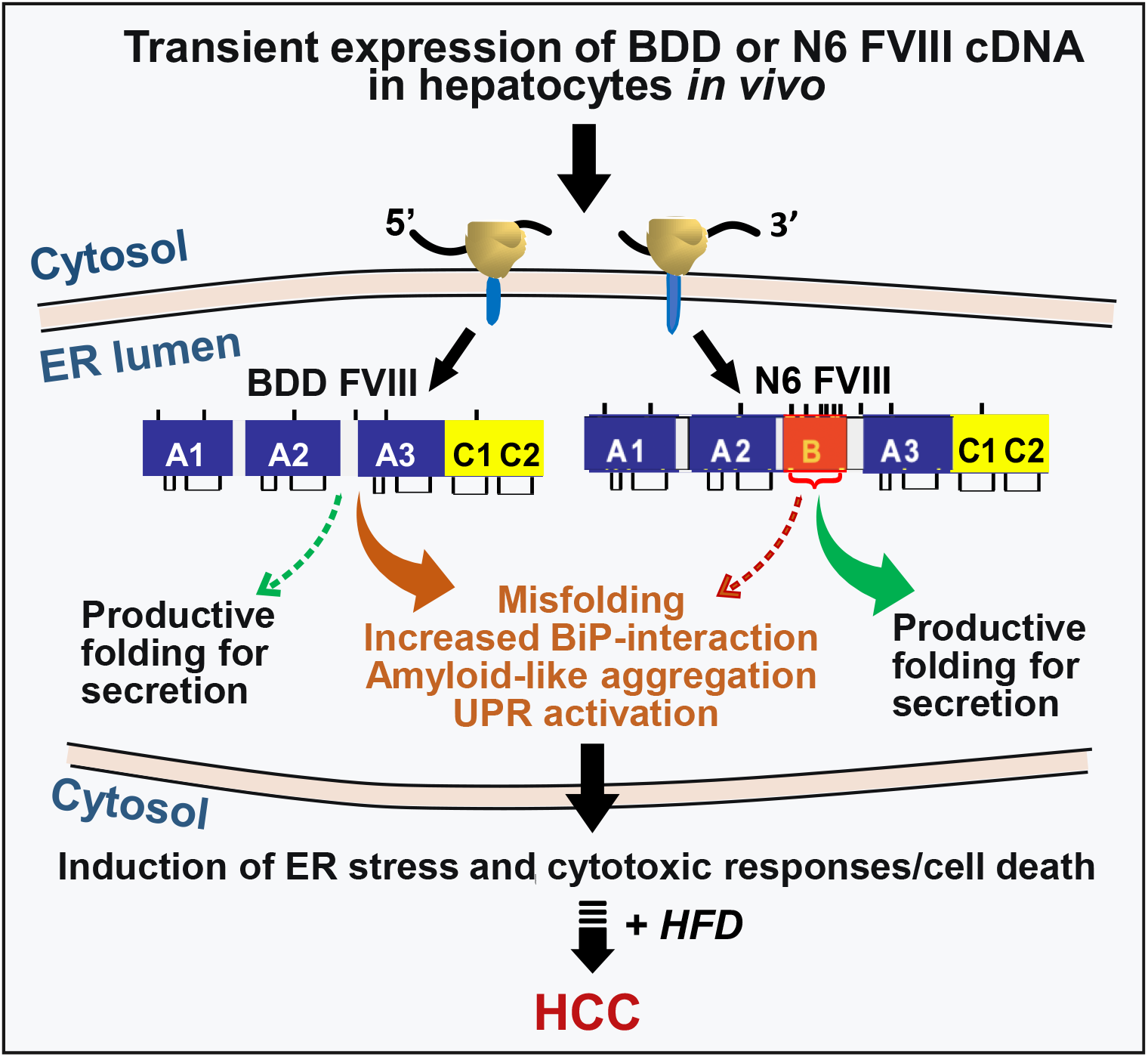

## Introduction

Hemophilia A (HA) is an X chromosome-linked bleeding disorder affecting 24.6 per 100,000 males at birth that results from deficiency of clotting factor VIII (FVIII).^1^ Protein replacement therapy with recombinant FVIII has significantly reduced morbidity and mortality associated with HA, although concerns remain. First, anti-FVIII inhibitory antibodies develop in ~25% of patients.^2^ Second, recombinant FVIII protein bears a high cost with limited availability due to low and variable production from mammalian host cells.^3^ FVIII secretion from mammalian cells is inefficient, partly due to FVIII protein misfolding, aggregation, and retention in the endoplasmic reticulum (ER).^4^ Misfolded FVIII activates the unfolded protein response (UPR) to resolve defective protein folding. However, upon chronic protein misfolding in the ER, the UPR shifts to an apoptotic program.

FVIII is a 330 kDa glycoprotein comprised of three major domains (A1-A2-B-A3-C1-C2).^5^ It is primarily produced by liver sinusoidal endothelial cells.^6^ The amino acid (aa) sequences in the A and C domains exhibit 40% amino acid identity between species, whereas the ~900 aa sequences within the large B domain show no homology other than a conserved large number (18) of N-linked oligosaccharides that are also observed in the B domains of FVIII from different species, as well as in the homologous clotting factor V. B domain-deleted FVIII (BDD),^7,8^ A1-A2-A3-C1-C2 (**Fig. 1**), is functional and similar to Refacto, used for protein replacement therapy for HA in the clinic (*i.e*., SQ-BDD FVIII; Pfizer).^9–11^ The only difference between BDD and SQ-BDD is that SQ-BDD contains an extra 14 amino acid linker (SFSQNPPVLKRHQR) at the junction between the A2 and A3 domains.^12,13^ Subsequent codon optimization improved SQ-BDD production leading to its use in ongoing HA gene therapy clinical studies.^12–16^ SQ-BDD is used for gene therapy via adeno-associated viral (AAV) vectors because the size of full-length FVIII is beyond the packaging limit of the vector genome, which is <5kb.^15^ However, BDD exhibits similar misfolding as intact FVIII, activates the UPR, and leads to hepatocyte death upon *in vivo* DNA vector delivery to hepatocytes in mice.^17^ Besides BDD, we described a partial B domain deletion molecule, herein (N6), that retains an additional 226 aa with 6 N-linked glycosylation sites^18^ (**Fig. 1**), and is secreted almost 10-fold more efficiently than full-length FVIII or BDD,^17^ presumably because the 6 N-glycans engage the intracellular lectin chaperone machinery, that was demonstrated to enhance FVIII trafficking from the ER to the Golgi compartment.^19–24^ Thus, N6 is secreted more efficiently and causes less UPR activation than BDD (**Fig. 1**).

**Figure 1.**
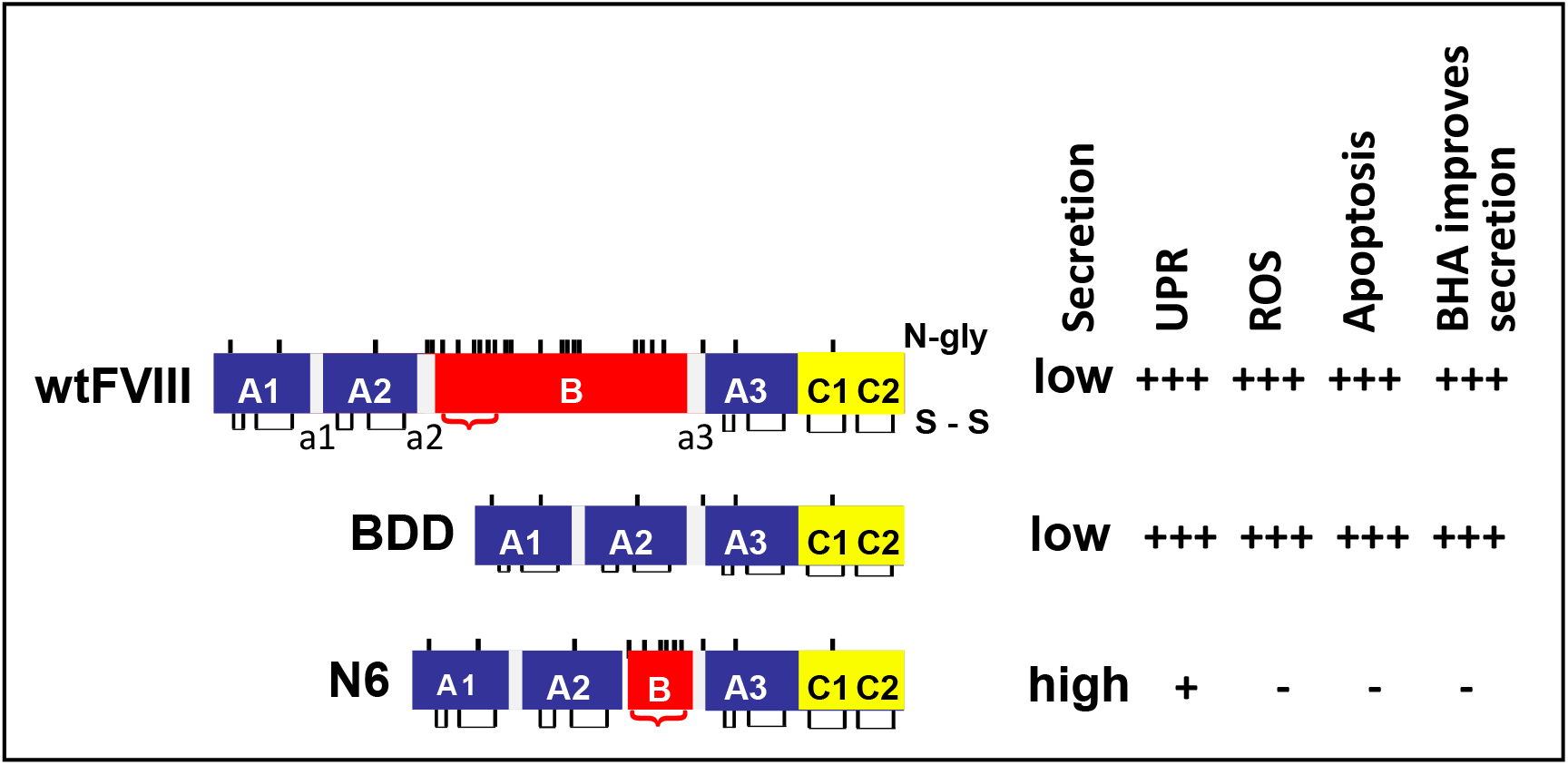
Hepatocyte responses to wtFVIII, BDD, and N6 upon hydrodynamic tail vein injection of vector DNA into mice. Relative FVIII secretion efficiency, activation of the UPR, production of reactive oxygen species (ROS), induction of apoptosis, and the ability for antioxidant butylated hydroxyanisole (BHA) to improve secretion were shown by us previously.^17^ a1-3=acidic regions. N-glycosylation sites (N-gly) and disulfide bonds (s-s) are depicted. Red bracket indicates residues from the B domain of wtFVIII that were retained in N6.

Presently, there are numerous ongoing liver-directed gene therapy clinical studies to deliver codon-optimized SQ-BDD via AAV to hepatocytes in men with severe HA. Initial results appeared to be very successful, with correction of FVIII levels into the normal range during the first year.^13,14^ Unfortunately, elevated liver enzyme levels were observed in the first year, and at a 5-yr^25^ follow-up, FVIII expression levels gradually declined to the lower end of the therapeutic range, starting in the second year.^26–29^ While immune response to the AAV vector is known to play an important role in liver pathology and the decline in FVIII expression in the clinical setting,^16,30^ it is possible that FVIII misfolding and UPR activation impact hepatocyte function, health, and/or survival. Therefore, there is an urgent need to understand mechanisms underlying inefficient FVIII secretion^26–28^ and the long-term pathologic effect of ectopic expression of FVIII in hepatocytes *in vivo*.

Previously, we demonstrated feeding a high fat diet (HFD) to mice that transiently over-express liver-specific urokinase-type plasminogen activator (uPA)-transgenic mice, which misfolds in the ER, initiates development of Non-Alcoholic Steatohepatitis (NASH) and hepatocellular carcinoma (HCC).^31^ Thus, it appears that either transient expression of a misfolded protein in the ER that activates the UPR followed by a HFD can initiate HCC development or there are unique properties of uPA that drive HCC progression. These findings led us to test whether transient hepatocyte expression of FVIII molecules with different folding efficiencies in the ER may also promote HCC in mice.

Here we studied the long-term outcome of transient expression of BDD, which aggregates in the cell and activates the UPR,^32^ and N6 that displays reduced aggregation and UPR activation.^17,18^ We delivered BDD and N6 expression vectors to murine livers by tail vein hydrodynamic DNA injection. These mice were then fed a HFD for 65 weeks starting at one week after vector DNA injection when FVIII expression was undetectable. We observed greater tumor formation in BDD vector injected mice than N6 vector injected mice. For the first time these findings support the notion that the degree of ER protein misfolding may be a significant factor in liver disease progression. Our findings point to a potential risk regarding ectopic expression of FVIII in hepatocytes in the context of HA gene therapy.

The potential for hepatic gene transfer to contribute to formation of HCC continues to be widely debated.^33,34^ For instance, a recent study found that AAV gene transfer to adult murine livers may cause HCC when animals receive a HFD.^35^ Hence, the combination of low-grade viral vector integration with the diet-induced hepatic steatosis and injury to the liver raises the risk of liver cancer formation. Herein, we found that even transient expression of a protein that is prone to misfolding and induction of ER stress (BDD-FVIII) in hepatocytes of adult mice is a trigger for HCC formation when followed by a HFD diet in the absence of viral delivery. Thus, this fundamentally new concept of protein expression-derived cellular stress predisposing to malignancy when followed by diet-induced inflammation is independent of molecular events caused by viral vectors. Importantly, the risk for HCC formation was reduced when a variant of FVIII was expressed that shows enhanced folding and secretion.

## Results

### FVIII forms aggregates that resolve at 1-2 hrs following energy repletion

Stable expression of BDD and N6 FVIII in CHO cells demonstrated that N6 expression was tolerated at an ~10-fold greater level than BDD.^18^ We previously characterized full-length FVIII aggregation by filtration of cell lysates through nitrocellulose (NC) membranes, which retain all cell proteins, and cellulose acetate (CA) membranes, which only retain proteins that have β-sheet aggregate structures.^32^ Lysate proteins retained on membranes were probed with FVIII or β-actin antibodies. Filtration through NC membranes demonstrated that intracellular steady state levels of N6 were ~5-fold greater than BDD, although secretion of N6 was significantly higher than BDD (**Fig. 2A**, red box). Filtration through CA membranes demonstrated significant aggregation of BDD and >4-fold less aggregation for N6 (ratio of CA/NC), although N6 was expressed at a ~5-fold greater level than BDD (**Fig. 2A**, blue box). Following energy depletion by treatment with 2-deoxyglucose (2-DG) and sodium azide (NaN_3_) to inhibit glycolysis and oxidative phosphorylation, respectively, FVIII aggregates accumulated in CHO cells that stably express wtFVIII (~500mU/ml/10^6^ cells/day, not shown),^32^ BDD (~1U/ml/10^6^ cells/day), and N6 (~10U/ml/10^6^ cells/day). We previously demonstrated that 2-DG, but not NaN_3_, was sufficient to induce wtFVIII aggregation.^32^ It is notable that NaN_3_ inhibition of oxidative phosphorylation is irreversible. After 2-DG removal and glucose replenishment, BDD and N6 aggregates began to disappear at 1-2 hrs, in a manner that did not require *de novo* protein synthesis as addition of the protein synthesis elongation inhibitor cycloheximide (CHX) did not alter aggregate dissolution or significantly reduce secretion of functional FVIII activity (**Fig. 2A**).

**Figure 2.**
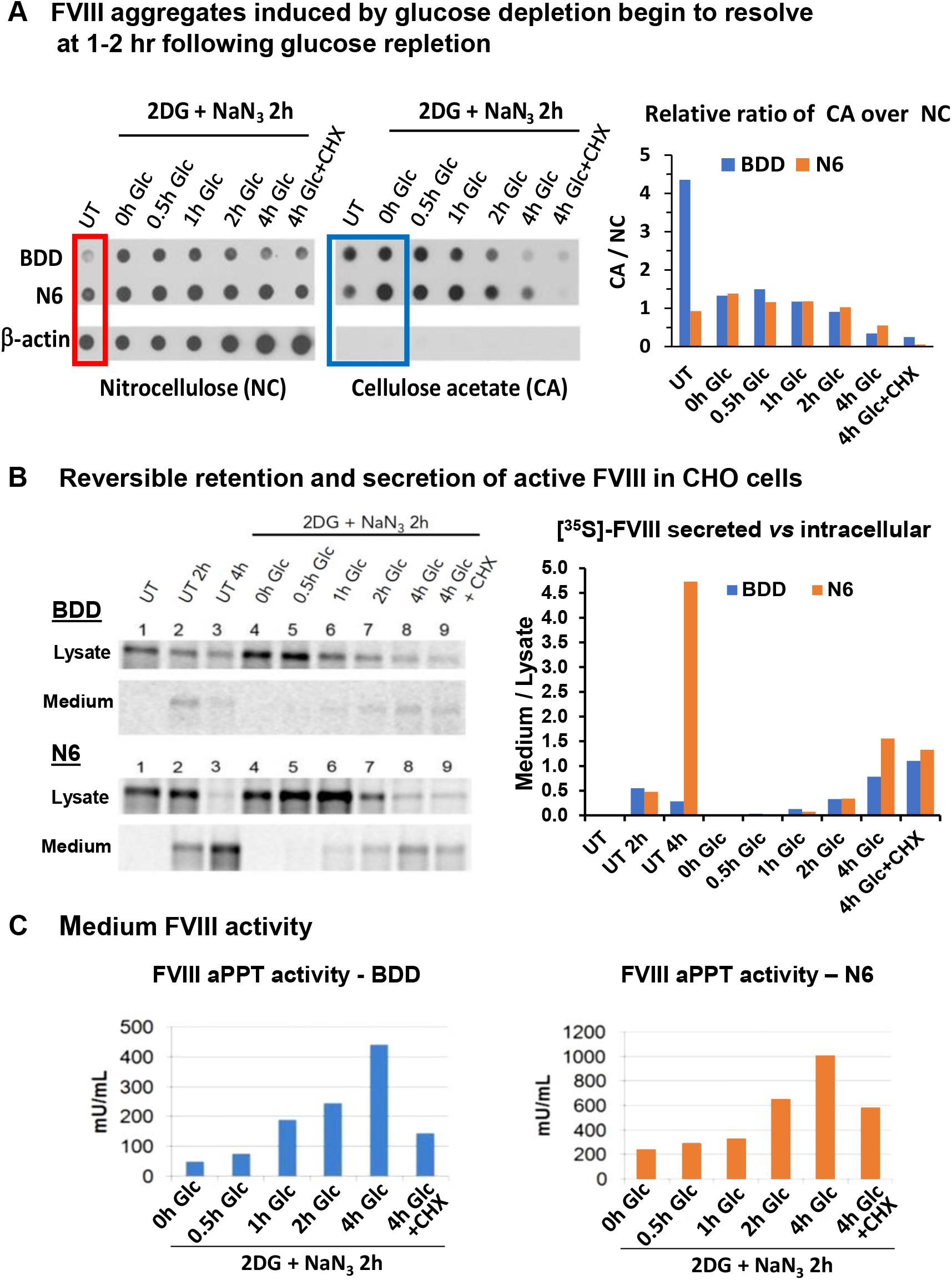
Reversible aggregation of BDD and N6 expressed in CHO cells. **A)** FVIII aggregates induced by glucose depletion begin to resolve at 1 hr following glucose repletion. CHO cells that express BDD or N6 were treated with either normal media or glucose-free media containing 10 mM 2DG and 20 mM sodium azide (2DG+NaN_3_). After 2 hr, cells were harvested or allowed to recover in complete media for the indicated times, or complete media with 10 μg/ml CHX. Cell lysates were filtered through nitrocellulose (left) or cellulose acetate (right) membranes and probed with FVIII or β-actin antibodies. Analysis of β-actin is from BDD expressing CHO cells. Red box = steady state levels of BDD and N6 in untreated cells. Blue box = aggregation of FVIII before and after 2DG + NaN_3_ treatment. **B)** Reversible retention and secretion of active FVIII in CHO cells. [^35^S]-Met/Cys pulse-chase CHO cells were treated in parallel as in Panel A. CHO cells were pulse-labeled for 20 min and then chased for 20 min with media containing excess unlabeled Met/Cys to complete synthesis of nascent chains (lane 1) before being treated with either normal media (lanes 2-3) or glucose-free media containing 10 mM 2DG and 20 mM NaN_3_. After 2 hr cells were harvested (lane 4) or allowed to recover in complete media for increasing times (lanes 5-8), or complete media with CHX (lane 9). Lysates and media were collected at indicated time points for FVIII IP and reducing SDS-PAGE. For FVIII aPPT activity assay of media, cells were treated in parallel, but not pulse-labeled. Lanes: 1: Untreated, 20’ chase; 2: Untreated 120’ chase; 3: Untreated 240’ chase; 4-9: 2DG for 120’; 5: 2-DG + 30‘ recovery; 6: 2-DG + 60’ recovery; 7: 2DG + 120’ recovery; 8: 2DG + 240’ recovery; 9: 2DG + 240’ recovery + CHX. **C)** FVIII activity in the medium via aPTT analysis of BDD and N6 expressed in CHO cells. All results (panels A-C) were from the same set of experiments performed in parallel.

To identify the destiny of aggregated BDD and N6, we performed [^35^S]-Met/Cys-pulse-chase labeling which confirmed that glucose deprivation retained BDD and N6 within the cell. Importantly, upon glucose repletion, intracellular aggregated BDD and N6 dissolved and were efficiently secreted into the medium (**Fig. 2B**). Analysis of FVIII activity in the conditioned media from cells treated under the same conditions as the pulse-chase assay demonstrated that the secreted FVIII is functional (**Fig. 2C**). Increasing amounts of functional FVIII appeared in the media as early as 1 hr following glucose repletion for BDD and at 2 hr for N6, which correlated with the rate of disappearance of intracellular aggregated FVIII. Although CHX treatment did not affect the amount of previously metabolically labeled FVIII secreted into the medium following glucose repletion, CHX treatment did reduce secreted FVIII activity to 30-60%, presumably due to inhibition of new FVIII synthesis. In sum, these results (**Figs. 2A-C**) indicate that a major percent of both BDD and N6 form metastable aggregates that can resolve to produce folded, functional, and secreted FVIII.

### Ectopic FVIII expression in hepatocytes causes protein aggregation in murine livers

Previously we demonstrated that full-length wtFVIII forms amyloid like fibrils upon expression in cultured cells.^32^ Hydrodynamic tail vein injection of vector DNA is an efficient method to express exogenous genes in hepatocytes.^18,36^ Using this technique, we confirmed expression of exogenous FVIII in hepatocytes of mice injected with BDD or N6 vector DNA induced the UPR, although the stress response to N6 expression was significantly attenuated.^17^ To determine if the stress response to FVIII expression in hepatocytes is associated with protein aggregation *in vivo*, we performed thioflavin-S (Thio-S) staining on the liver sections to specifically identify amyloid-like protein aggregates. The livers of BDD vector-injected mice showed significantly more Thio-S positivity than N6 vector-injected mice, evidenced by colocalization of Thio-S staining with FVIII immunostaining (**Fig. 3**). Similar to BDD expression in CHO cells,^32^ these findings demonstrate that ectopic expression of BDD in hepatocytes *in vivo* also leads to intracellular accumulation of amyloid-like FVIII aggregates in the liver. As expected, neither FVIII nor Thio-S stains were observed in mice that received the parental vector pMT expressing the cytosolic well-folded enzyme dihydrofolate reductase (DHFR) (**Fig. 3**).

**Figure 3:**
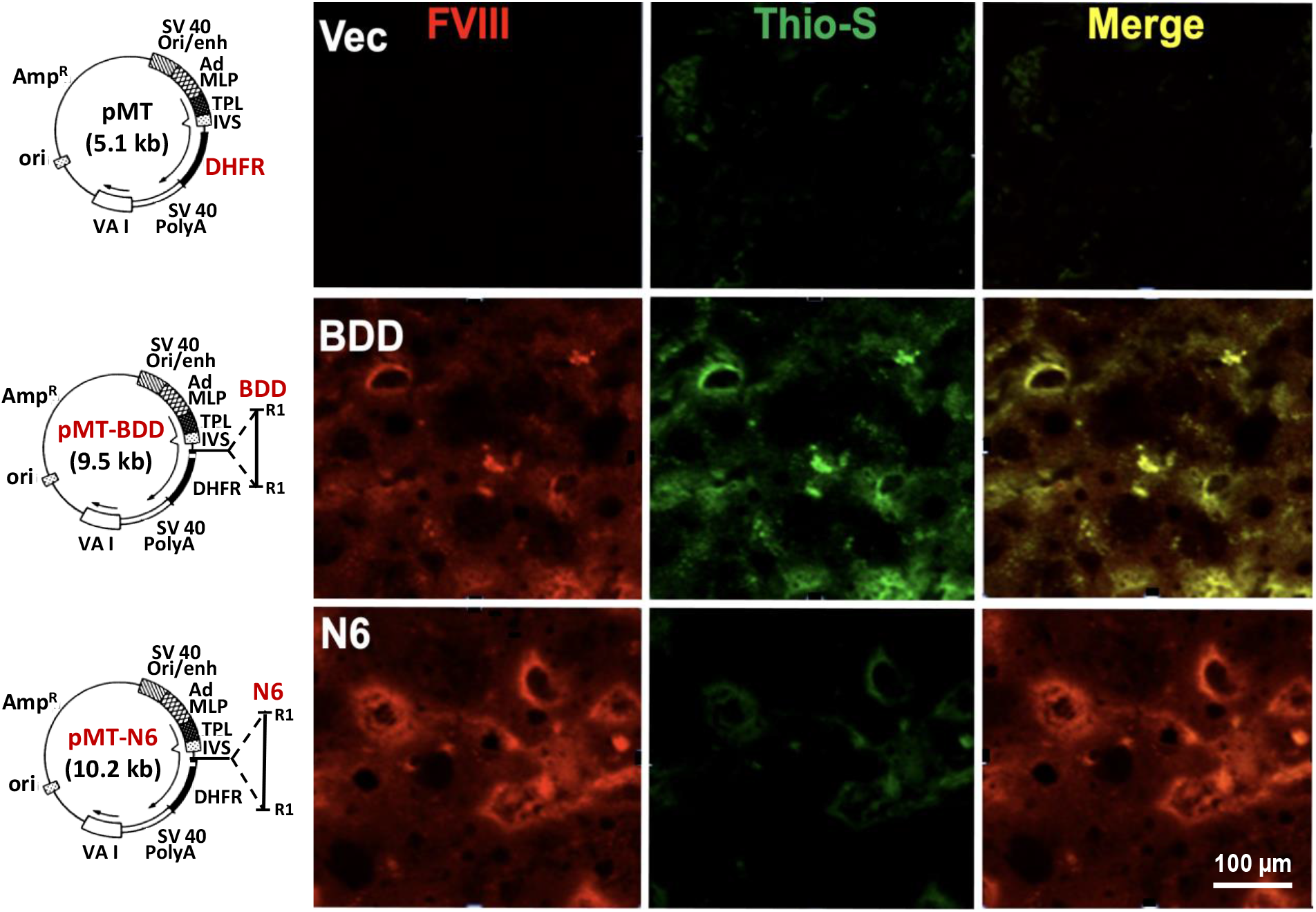
BDD expression in hepatocytes *in vivo* causes accumulation of amyloid-like aggregates that colocalize with FVIII in liver sections. Liver sections were prepared from mice injected with the indicated vector DNAs as described^17^ and stained for FVIII (red) and Thio-S (green).^32^ Shown are representative images from these analyses. Details of the vector constructs are depicted with plasmid maps on the left-hand side. The parental vector pMT (^69^, Addgene) contains a transcription unit that is composed of the SV40 origin and enhancer element from the early promoter, the adenovirus major late promoter containing the majority of the tripartite leader present in adenovirus late mRNAs, a hybrid intron (IVS), the cytosolic dihydrofolate reductase (DHFR)-coding region, and the SV40 early polyadenylation signal (SV40 PolyA). The parental vector efficiently expresses DHFR.^69^ Human BDD or N6 coding sequences were inserted upstream of the DHFR-coding region at the *Eco*R1 restriction sites (R1).

### BDD misfolds in murine hepatocytes *in vivo*

To provide mechanistic insight into BDD misfolding and UPR activation, we characterized BDD and N6 interaction with the ER chaperone BiP/GRP78. Misfolded proteins bind BiP whereas well-folded proteins do not.^37,38^ Therefore, protein interaction with BiP can be used as a surrogate to measure ER protein misfolding. However, there are no available antibodies that can quantitatively co-immunoprecipitate (IP) BiP-client protein complexes. Hence, we created a genetically modified mouse with a 3x FLAG tag inserted into the endogenous BiP/GRP78/*Hspa5* locus (*BiP-Flag* mice) for quantitative isolation of BiP-client protein complexes using anti-FLAG magnetic beads,^39^ Tagging BiP with 3X FLAG in this manner did not alter BiP function or BiP expression, which is essential to measure physiological interactions **(Figs. 4A-B**,^39,40^). Importantly, hepatocyte-specific BiP-Flag displays the same expression level and is regulated identically as the untagged wildtype BiP (**Fig. 4B**). To elucidate the folding status BDD and N6 ectopically expressed in murine hepatocytes, we expressed BDD and N6 in hepatocytes of *BiP-Flag* homozygous (*BiP-Flag^+/+^*) mice by hydrodynamic tail vein delivery of vector DNA and analyzed the interactions between BiP-Flag and BDD or N6 through anti-FLAG co-IP of BDD and N6 (**Fig. 4C**). Consistent with our previous observations^17^, hepatic BDD levels in BDD vector-injected mice were significantly higher than those in the N6 vector-injected mice (**Fig. 4C**, lanes 5-10 vs lanes 11-16 and **Fig. 4D**, left panel), probably due to the greater retention of BDD in the ER. Importantly, anti-FLAG IP pulled down a significant portion of BDD from the livers of BDD vector-injected *BiP-Flag^+/+^* mice (25%; **Fig. 4C**, lanes 24-26 vs 8-10 and **Fig. 4D**, right panel), while N6 was pulled down at significantly lesser level (16%; **Fig. 4C**, lanes 24-26 vs 30-32 vs 14-16 **Fig. 4D**, right panel). These results support the notion that BDD exhibits greater misfolding than N6 in hepatocytes of murine livers, which likely accounts for the different degrees of UPR induction.^17^

**Figure 4.**
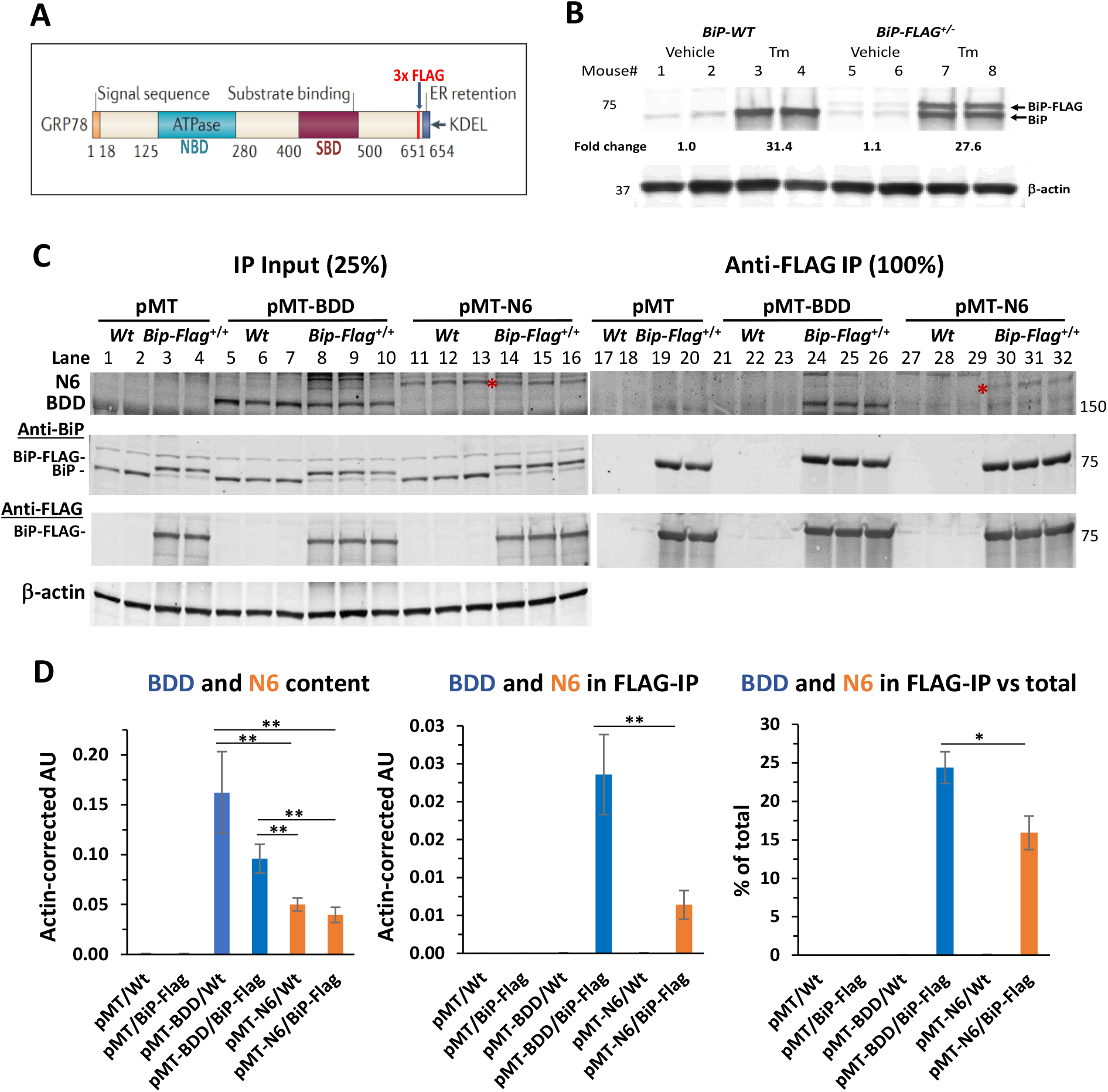
Epitope-tagging of the endogenous murine BIP/GRP78/*Hspa5* locus demonstrates misfolding of BDD *in vivo*. **A)** A diagram illustrating location of insertion of 3x Flag tag to C-terminal region of BiP; **B)** Western blot demonstrating similar changes in hepatic levels of endogenous wtBiP (BiP) and BiP-Flag in mice heterozygous for *BiP-Flag* allele (10 days after activation of BiP-Flag expression in the liver) in response to treatment with tunicamycin for 24 hr. **C)** Co-IP of BDD (lanes24-26) and N6 (lanes30-32) from liver lysates of pMT-BDD or pMT-N6-injected *BiP-Flag* homozygous mice (*BiP-Flag^+/+^*) with anti-FLAG magnetic beads. Hepatic level of BDD and N6 were determined by direct Western blot on liver lysates (lanes 5-10 and lanes 11-16 for BDD and N6, respectively). Red asterisk = N6 FVIII bands. **D)** Quantification of BDD and N6 bands shown in panel C. Data are represented as mean ± S.E.M. * P<0.05, ** P<0.001. Each lane in the images for Western blots represents an individual mouse, i.e., the number of mice (n) in each group: n=2 for pMT/Wt and pMT/BiP-Flag; n=3 for pMT-BDD/Wt, pMT-BDD/BiP-Flag, pMT-N6/Wt, and pMT-N6/BiP-Flag, respectively.

### Lentiviral transduction of BDD into human primary hepatocytes activates the UPR

We next asked whether BDD expression can activate the UPR in human hepatocytes. We used mouse-passaged human primary hepatocytes (mpPHH)^41^ to investigate the cellular response to ectopic BDD expression. The mpPHH were transduced *in vitro* with increasing amounts of a lentiviral vector that expresses a codon-optimized SQ-BDD cDNA^12^ that is used in all current liver-directed HA gene therapy trials.^13,14,16^ Western blot analysis demonstrated SQ-BDD expression induced UPR markers BiP, eIF2α phosphorylation, and CHOP, despite its weak signal, at 10 days after infection (**Fig. 5**). Interestingly, the protein levels of two UPR-induced genes in the liver, hepcidin^42^ and cysteine rich with EGF like domains 2 (CRELD2), an oncogene associated with HCC that promotes survival upon ER stress,^43–46^ were increased in BDD-expressing mpPHH (**Fig. 5)**. These results for the first time show that ectopic BDD expression induces ER stress in <u>human</u> hepatocytes.

**Figure 5.**
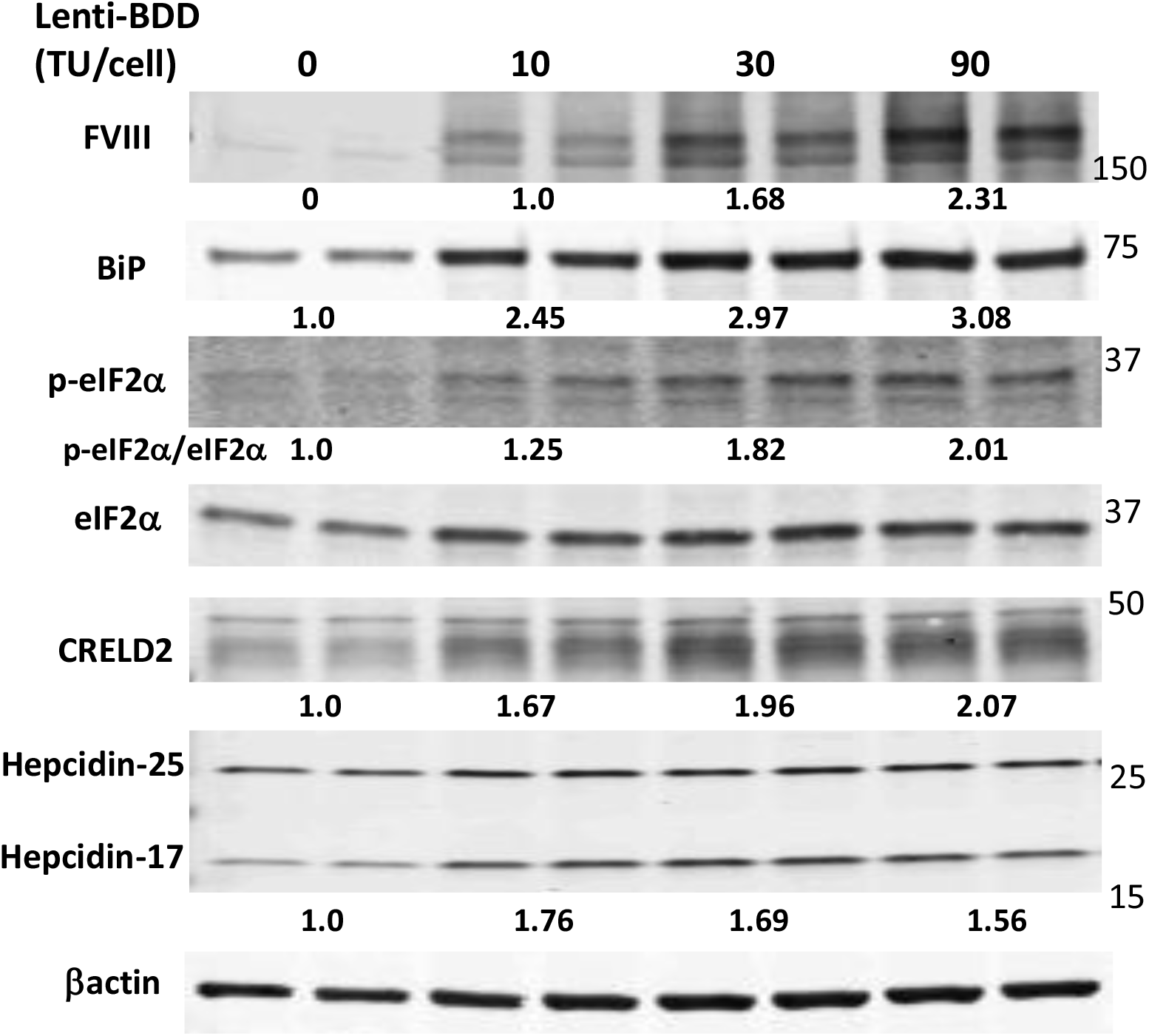
BDD expression induces the UPR in mouse-passaged primary human hepatocytes (mpPHH). mpPHH were transduced with a lentiviral vector expressing a codon-optimized form of SQ-BDD (Lenti-BDD) at the indicated vector doses and cultured in a hepatocyte-defined medium. The transduced mpPHH were harvested for Western blot analysis 10 days post-transduction. Each lane represents individual culture well. The numerical numbers represent averaged fold changes after correction with loading control *β*-actin. Experiment was repeated twice with different batches of mpPHH and similar results were obtained.

### HCC development correlates with the degree of ER protein misfolding

Previous findings suggest protein misfolding in the ER can initiate NASH and HCC development^31^. Therefore, we tested whether two different human FVIII variants that misfold to different degrees can initiate HCC development. *C57BL/6J* mice were subjected to hydrodynamic injection of parental vector pMT, pMT-BDD, or pMT-N6 vector DNA.^17^ After 24 hr, plasma levels of N6 FVIII were significantly higher than those for BDD FVIII (**Figs. 6A** and **S1A**), consistent with our previous observations.^17^ After 1 wk, when FVIII expression was not detectable (**Figs. S1B-D**), mice were fed a 60% HFD for 65 wks and then analyzed for liver surface tumors. No tumors, neither adenomas nor adenocarcinomas, were observed in mice that received parental vector pMT DNA that expressed the cytosolic DHFR. In contrast, all mice that received pMT-BDD expression vector developed tumors, while reduced tumor formation was observed in mice that received pMT-N6 vector (**Fig. 6B**). All mice in the pMT-BDD group had 1 liver surface tumor whereas a couple tumors were observed in 3 mice from the pMT-N6 group.

**Figure 6.**
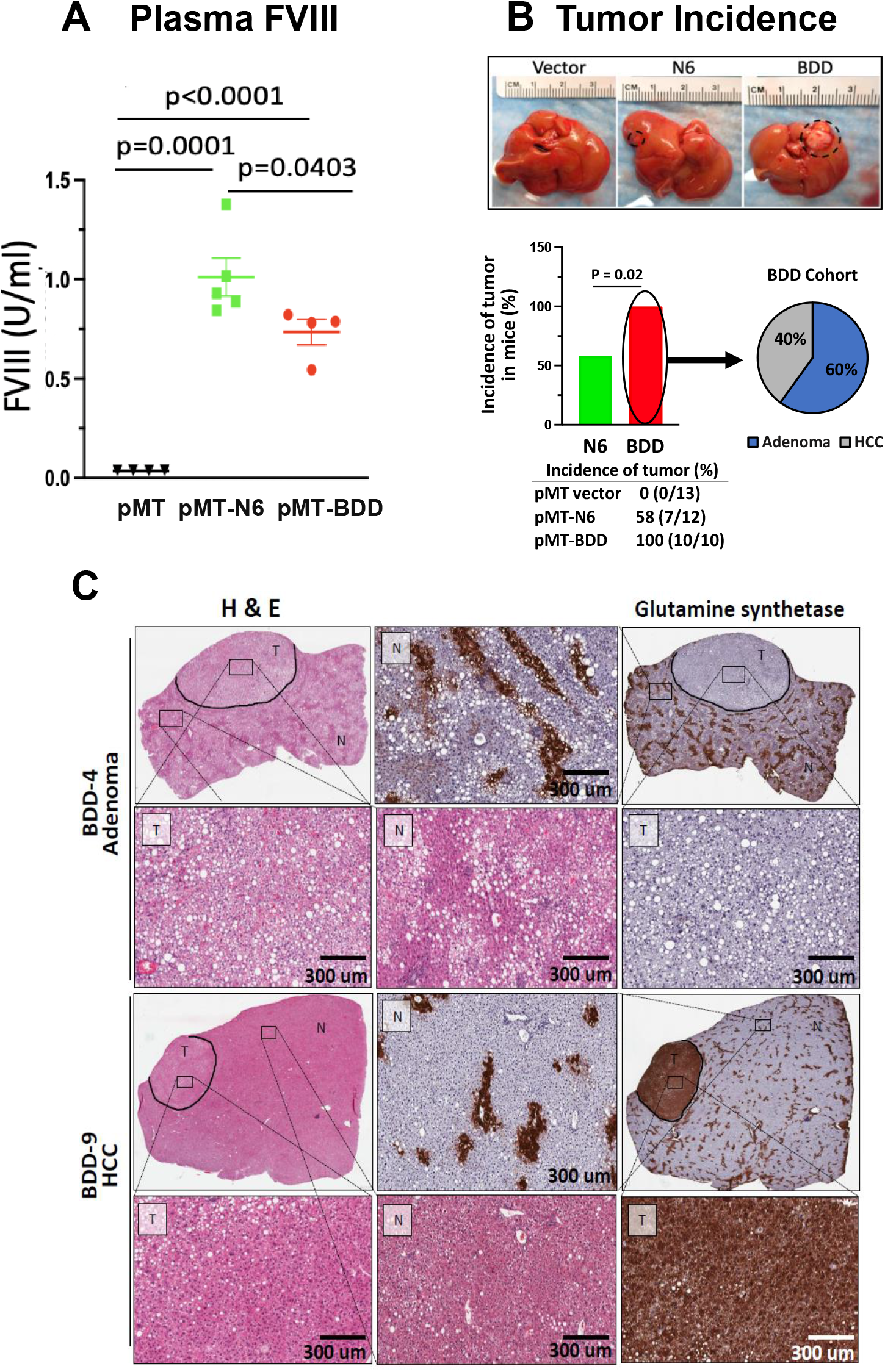
Transient FVIII expression in hepatocytes causes liver tumors in mice in the context of high-fat diet (HFD) feeding. Wild type *C57BL/6J* male mice were injected with pMT vector, pMT-N6 or pMT-BDD DNA vectors by hydrodynamic tail vein injection. **A)** Plasma FVIII levels at 24 hr after vector DNA injection. Data shown are Mean ± SE (N=4 for pMT and pMT-BDD, N=5 for pMT-N6). **B)** Incidence of liver tumors in pMT-BDD- or pMT-N6 injected mice after a 65-wk-HFD treatment. The difference between the two groups (*ie*., pMT-N6 & pMT-BDD) were tested for statistical significance using the Chi-square test (degrees of freedom=1, 95% confidence intervals). **C)** HCC but not adenoma exhibited positivity for glutamine synthase staining. Both tumor tissue (T) and non-tumor tissue (N) are shown in the high magnification images.

The tumors (both adenomas and carcinomas) in the above mice were evaluated and extensively characterized independently by three clinical pathologists. A detailed summary of this histopathological analysis is provided in **Tables SI–S3**. Firstly, reticulin staining is intended to demonstrate reticular fibers surrounding tumor cells. Tumors that retain a reticulin network are generally benign or pre-neoplastic, whereas HCC loses the reticulin fiber normal architecture. Secondly, immunohistochemistry staining of CD34, glutamine synthetase (GS), β-catenin, glypican 3, and CD44 were also performed to differentiate the adenomatoid dysplastic nodule from the HCC lesions. Although no tumors were detected in any of the 13 mice injected with parental vector pMT, we observed that 58% (7/12) of mice that received pMT-N6 vector developed liver tumors; in contrast to 100% (10/10) of mice that received pMT-BDD vector (**Fig. 6B**, p<0.02, Chi-square test). The findings suggest a higher penetrance of tumors in the presence of BDD misfolded protein. Detailed histopathological assessment of the pMT-BDD-injected mice demonstrated that 6 of the 10 lesions were adenomas and 4 carcinomas (**Fig. 6B**), while the pMT-N6-injected mice developed 4 adenomas and 3 carcinomas (**Fig. S2A**). CD34 demonstrates neovascularization in HCC, while nontumorous hepatic sinusoids do not stain with CD34.^47^ The adenomas were overall negative for CD34 whereas the carcinomas displayed a patchy strong CD34-positive staining (**Fig. S3**), further confirming the malignancy.^47,48^ Glutamine synthetase stains positively for zone 3 of the liver parenchyma. Most carcinomas had positive GS staining (**Fig. 6C)**, indicative of an early-stage HCC.^49^ However, a few well-differentiated HCCs and an early evolving HCC, arising from adenomas, were GS negative. As expected, β-catenin immunostaining showed identical results to GS as they are both frequently overexpressed in early HCC.^49^ Finally, the adenomas in the BDD cohort were positive for CD44, with the sinusoidal lymphocytes staining while the adenomas in the N6 cohort were negative (**Fig. S3**). Based on these histopathological assessments, the distribution of adenomas and carcinomas appears to be similar in both groups (**Figs. 6C** and **S2A**). However, BDD mice developed more tumors and thus a more aggressive phenotype. In addition, the staining intensity and cellular distribution of the tumor makers differed considerably in the liver tumors observed in our mice (**Table S3**), indicating that these tumors were at different stages of development towards HCC. It is important to note that none of the tumors found in these mice expressed BDD or N6, as demonstrated by the analysis of DNA isolated from the liver tumors and their adjacent non-tumor tissues collected at the study end point (**Table S4)**, providing evidence supporting the notion that transient expression of misfolded FVIII, over the first week after DNA delivery, in the hepatocyte combined with HFD feeding is sufficient to induce HCC.

## Discussion

Hepatic gene transfer of codon-optimized SQ-BDD FVIII cDNA using AAV vectors has emerged as a promising therapeutic approach for HA.^13,14^ Substantial improvement in hemostasis was documented at a 5-yr follow-up,^25^ although the approach has encountered two key hurdles: 1) A requirement for very high vector doses to drive FVIII expression in hepatocytes; and 2) In addition to signs of liver damage, transgene expression declined over time. ^26–29^ Both may be directly linked to FVIII misfolding and retention in the ER. Durability of transgene expression is a major factor for HA gene therapy as it is safely and effectively treated by prophylaxis with recombinant FVIII as well as FVIII-bypassing molecules such as a bispecific antibody^29^ raising the safety and efficacy bars for HA gene therapy regimens. The significance of this problem was recently highlighted.^26,50,51^ Here, we demonstrate that cellular stress resulting from transient expression of FVIII in hepatocytes may contribute to liver tumorigenesis. Encouragingly, we find that a more efficiently secreted FVIII variant can be developed to reduce this risk. While there are differences in the biology of human and murine HCC and AAV gene transfer differs from plasmid vectors, these outcomes warrant further investigations into the potential for gene therapy approaches to cause tumors in patients with HA.

Our previous work identified a FVIII derivative, N6, that is more efficiently secreted and less prone to aggregation than present B-domain deletion molecules used for HA gene therapy.^17,18^ In addition to the increased secretion efficiency of N6 compared to BDD, N6 elicits reduced cellular toxicity, as measured by UPR activation, apoptosis, levels of reactive oxygen species (ROS), protection by antioxidant BHA treatment, and HCC (**Fig. 1**).^17,18,52,53^ Using mice with a FLAG tag inserted into the endogenous *BiP/Hspa5* locus for quantitative FLAG IP of BiP-client protein complexes,^39^ we found that BDD displays greater interaction with BiP compared to N6, *i.e*., a quarter of intrahepatocellular BDD binds BiP (**Fig. 3D**), indicative of misfolding. The *BiP-Flag* mouse provides the ability to evaluate the misfolding propensity of FVIII derivatives considered for gene therapy. Ectopic expression of BDD produces a higher level of misfolded FVIII, which can induce a series of cytotoxic responses, including activation of the UPR,^17^ ultimately leading to initiation of HCC development.

Our findings suggest that characterization of the folding efficiency, host response, and safety in model organisms and non-human primates is essential to ensure safety over the lifetime of an individual, given that HCC takes years to develop in humans. A case of HCC did occur in a hemophilia B patient after hepatic AAV-mediated factor IX gene transfer^54^ which was unlikely caused by gene therapy as it occurred early after gene transfer and the patient had multiple risk factors that pre-disposed to HCC. While HCC formation was observed in murine studies with AAV vectors using other transgenes,^35^ this was attributed to insertional mutagenesis and the relevance of this observation to the safety of HA gene therapy remains to be understood. However, our new results show that high levels of FVIII gene expression by itself, even if transient and very likely independent of insertional events as our studies did not use viral vectors, can induce HCC in the context of a HFD. All murine models of NASH and HCC progression require a hypercaloric diet, presumably to induce an inflammatory environment.^55,56^ HCC has not been observed in animals treated with AAV-FVIII vectors, but these animals were not subjected a HFD challenge. Interestingly, neonatal AAV gene transfer (expressing other transgenes) led to HCC under a normal dietary regimen but transduction of the same AAV vectors to adult mice did not induce HCC unless they were fed a HFD leading to NAFLD,^35^ which was associated with increased inflammation and hepatocyte proliferation, consistent with our findings. Nevertheless, it should be considered that our studies do not reflect the course of liver disease in humans, nor the therapeutic approach. For instance, we used DNA delivery in our mouse studies because we previously characterized detailed responses to BDD vs N6.^17^ It is of great importance to investigate if the amount of FVIII produced by hepatocytes under a setting of AAV gene transfer reaches a threshold required to set off the cascade of molecular events that promotes HCC. Long-term follow-up in patients who have high levels of FVIII expression will be required to answer this question.

Our studies also support the need for careful monitoring of liver function over the time course of gene therapy treatment. We demonstrate the role of diet when expressing a therapeutic protein in the liver that is prone to misfolding and ER stress. While AAV-FVIII gene transfer has not been found to cause the level of hepatocyte apoptosis and liver injury that may occur with DNA vectors, multiple studies have shown induction of ER stress markers in mice in response to AAV-FVIII,^57–59^ and liver injury (albeit of unknown origin) occurred in patients treated with high-dose AAV-FVIII.^13,14^ We now directly show that expression of SQ-BDD in primary human hepatocytes induces ER stress and the UPR in the dose-dependent manner (**Fig. 5**). Together, our study provides experimental evidence demonstrating the need for rigorous scientific investigation towards the pathophysiological consequences upon AAV-mediated FVIII expression in hepatocytes. HCC is the most common primary liver cancer and the 3rd leading cause of cancer deaths worldwide.^60^ The incidence of HCC has tripled since 1980 and is the most increasing cause of cancer mortality in the U.S.^61^ While the incidence of viral-related HCC is declining, HCC related to metabolic stress is on the rise, due to the global increase in obesity and the associated metabolic syndrome.^62^ It is currently estimated that NASH, which is already the leading cause of liver transplants in developed countries, will become the dominant HCC etiology. A quarter of the world’s population suffers from non-alcoholic fatty liver disease (NAFLD) characterized by abnormal lipid accumulation in hepatocytes. Approximately ~20% of NAFLD patients eventually develops NASH with inflammation, hepatocyte ballooning, Mallory Denk Bodies and cell death, a subset of which further develop fibrosis, cirrhosis, and HCC.^63^ Both ER stress and activation of the UPR are documented in many different human diseases,^64,65^ including NASH and HCC.^66–68^ The prevalence of NASH in the general population may increase the risk for HCC associated with the hepatocyte-targeted gene therapy.

In conclusion, we provide the first evidence that even transient hepatic expression of a protein that is prone to misfolding in the ER to induce the UPR, can trigger HCC formation when followed by a second insult to the liver such as a HFD diet. While these findings in mice using DNA vectors do not necessarily mean that humans treated with viral vectors in current clinical approaches are at increased risk of cancer formation, limited levels of expression of such proteins in hepatocytes and/or use of better secreted variants should be considered in clinical development. Importantly our findings demonstrate that a factor, independent of viral toxicity, can be contributed by protein misfolding in the ER.

## Materials and Methods

### Cell lines

Parental Chinese hamster ovary cells (CHO-K1) and 2 CHO-K1 clones were engineered for constitutive BDD (CHO-BDD) or N6 (CHO-N6) expression were previously described ^18^.

### Reagents and Standard Methods

All reagents and standard methods are specified in Supplemental Materials.

### Statistical Analysis

Statistical analyses were performed using GraphPad Prism 9 and a two-tailed Student’s *t*-test. Chi-square test was used for comparing of tumor incidence between pMT-BDD and pMT-N6 groups. A p value < 0.05 was regarded as statistically significant (p < 0.05 *, p < 0.01 **).

## Acknowledgments

This work was supported by NIH grants CA198103 and DK113171 to R.J.K. and in part by the NIDDK-funded San Diego Digestive Diseases Research Center (P30DK 120515). Portions of this work were supported by NIH 1P01HL160472-01 (to R.W.H., R.J.K. and Y.P.D.J.). We thank Takeda Pharmaceuticals U.S.A., Inc., for providing us with the Sepharose-coupled and uncoupled monoclonal anti-human FVIII antibodies for this work.

## Author Contributions

R.J.K. conceived the project, designed, and interpreted experiments, and wrote the manuscript with Z.C. and A. K. L. A.K.L., Z.C., R.D., C.L., L.G., J.M., J.Y. and C.Z. performed various *in vivo* and *in vitro* experiments. Z.G., A.L. and P.M. provided expert consultation and supervision on tumor analyses. Y.P.D.J. and R.W.H. interpreted data and edited the manuscript. R.J.K. supported the study.

## SUPPLEMENTAL MATERIALS

### Materials and Methods

#### Materials for cellular and molecular biology

Anti-factor VIII heavy chain monoclonal antibody coupled to Sepharose CL-4B was a kind gift from Baxter/Baxalta Corp. FVIII-deficient and normal pooled human plasmas were obtained from George King Biomedical (Overland Park, KS). Activated partial thromboplastin (automated aPTT reagent) and CaCl2 were purchased from General Diagnostics Organon Teknika (Durham, NC). Dulbecco’s modified Eagle medium (DMEM), glucose-free DMEM, alpha-modified essential medium (alpha-MEM), cysteine/methionine-free DMEM, and fetal bovine serum (FBS) were purchased from Gibco BRL. FVIII:C-EIA was purchased from Affinity Biologicals. Anti-β-actin antibody, 3-methyladenine, 2-deoxy-D-glucose (2-DG), and sodium azide (NaN_3_) were obtained from Sigma Aldrich. Anti-FVIII antibody (GMA012) was obtained from Green Mountain. [^35^S]-Methionine/Cysteine was obtained from MP Biologicals. Mouse and rabbit horseradish peroxidase conjugated secondary antibodies, Prolong Antifade Gold and Complete Protease Inhibitor Cocktail were obtained from Promega. Supersignal West Pico ECL was obtained from Thermo. Mouse FAB fragments, Dylight 549 conjugated anti-mouse fab fragments and Texas-Red conjugated anti-mouse secondary were obtained from Jackson Immunoresearch (West grove, PA).

#### Mice

Male *C57BL/6J* mice were purchased from Jackson Laboratory and maintained at the Sanford-Burnham-Prebys Medical Discovery Institute animal facility. Mice were euthanized by CO2 inhalation for liver harvest. All animal protocols were reviewed and approved by the Institutional Animal Care and Use Committee at the SBP Medical Discovery Institute.

#### Hydrodynamic tail vein injection of plasmid DNA and HCC development

The expression vectors for BDD and N6 were previously described.^1^ Six wk-old male mice were used for these experiments. Hydrodynamic tail vein DNA injection was performed according to previous publications.^2,3^ In summary, 100μg plasmid DNA was diluted in 2.5ml saline and injected into mice through the tail vein. One wk after the injection, mice were fed 60% HFD for 65 wks after which their livers were harvested for tumor evaluation.

#### Glucose depletion and repletion

For glucose depletion, cells were treated with ATP-depleting medium (glucose-free DMEM containing 20 mM 2-DG and 10 mM NaN_3_) for 2 hr. To replete glucose, fresh media was replaced for the indicated time. Cycloheximide (CHX) at a final concentration of 10 μg/mL was added to the repletion media where indicated.

#### Factor VIII activity and antigen analysis

FVIII activity was measured by a 1-stage activated thromboplastin time (aPTT) clotting assay on an MLA Electra 750 fibrinometer (Medical Laboratory Automation, Pleasantville, NY) by reconstitution of human FVIII-deficient plasma. The FVIII plasma standard was FACT plasma (normal pooled plasma) from George King Biomedical. FVIII antigen was quantified by an anti-FVIII sandwich enzyme-linked immunosorbent assay (ELISA) method using the Affinity Biologicals FVIII:C-EIA commercial kit (Affinity Biologicals Inc., Hamilton, ON, Canada) according to the manufacturers’ instructions.

#### Metabolic labeling

Cells were subcultured 24 hr prior to labeling in 60 mm dishes (approximately 10^6^ cells/plate) and were 80% confluent at the time of labeling. Cells were washed twice in cys/met free DMEM and incubated in cys/met-free DMEM for 10 min prior to labeling. Cells were labeled in 0.5 ml Cys/Met free DMEM containing 100 μCi/mL (BDD-FVIII and N6-FVIII) for 20 min and chased for the indicated times with conditioned medium (either 2-DG and NaN_3_-containing medium or normal medium) containing excess unlabeled Cys and Met and 10 μg/ml aprotinin. For glucose-depletion/repletion conditions, depleting medium was removed after 2 hr and replaced with normal medium containing excess unlabeled Cys and Met and aprotinin as above. At the end of the chase period, conditioned media was collected. Cells were rinsed three times in phosphate buffered saline (PBS) and harvested in 1 ml lysis buffer [50 mM Tris-HCl, pH 7.4, 150 mM NaCl, 0.1% (v/v) Triton X-100, and 1% (v/v) IGEPAL] containing Complete Protease Inhibitor Cocktail and 1 mM phenylmethylsulfonyl fluoride. Lysates were incubated on ice 30 min, followed by centrifugation at 15,000 x g for 10 min. Post-nuclear supernatant was then assayed for protein content using BCA assay (Bio-Rad). Equal amounts of protein and corresponding amounts of media were subjected to FVIII immunoprecipitation (IP) using anti-FVIII coupled Sepharose CL-4B beads at 4°C overnight. IPs were washed 4 times with lysis buffer and proteins were separated by reducing SDS-PAGE. IP’d proteins were visualized by autoradiography and band intensities were quantified using ImageQuant normal media for indicated times. CHX at a final concentration of 10 μg/mL was added to the repletion media where indicated.

#### Interaction of BDD and N6 with BiP *in vivo*

The *BiP-Flag* mice were generated as described.^4^ Mice were fed a regular mouse chow diet. Induction of BiP-Flag expression was initiated at the age of 6 wks by transduction with AAV8-TGB-Cre.^4^ AAV8-TGB-Cre-transdcued wild type littermates were used as controls. At 6 wks after AAV8-TGB-Cre-transduction, the mice received the parental pMT2 vector, BDD vector, or N6 vector as detailed in Figure 3 through hydrodynamic tail vein DNA injection^3^ and were sacrificed to collect liver tissues 26 hr post-vector DNA injection.

Liver samples were harvested in lysis buffer [0.15 mM NaCl, 0.5% Triton X-100, 20 mM HEPES, pH 7.4, 1x Protease inhibitor cocktail (Fisher)]. The liver lysates were centrifuged at 10K x g for 10 min and the resultant supernatants were used for subsequent analyses. For anti-Flag IP, 120 μg of lysate proteins in 300 μl of lysis buffer was mixed with 15 μl of M2 anti-Flag magnetic beads (Sigma) and incubated with rotation at 4°C for 4 hr. At the end of this incubation, the anti-Flag beads were washed 3X with ice-cold lysis buffer (1 ml each). Proteins in the washed anti-Flag and anti-FVIII beads were eluted with SDS-PAGE loading buffer, separated by SDS-PAGE under reducing conditions and transferred onto Nitrocellulose membranes for Western blotting along with 32 μg proteins for each liver lysate sample used for Flag IP (ie., 25% of the input). A mouse anti-human FVIII monoclonal antibody (GAM012, Santa Cruz Biotechnology) was used for BDD and N6 detection.

#### *Ex vivo* studies with mouse-passaged primary human hepatocytes

Mouse-passed primary human hepatocytes (PHH) were prepared and cultured as described.^5^ They were transduced with a lentiviral vector expressing a codon-optimized form of SQ-BDD coding sequence (Lenti-BDD)^6^ at the indicated doses 3 days after isolation and being cultured *ex vivo* and harvested for Western blot analysis 10 days after transduction.

Antibodies used were as follows: Rabbit anti-BiP, rabbit anti-phospho eIF2α and rabbit anti-eIF2α were from Cell Signaling Technology, (Danvers, MA). Mouse anti-human FVIII was from Green Mountain Antibodies (Burlington, VT); Rabbit anti-CRELD2 and mouse anti-hepcidin antibodies were from Proteintech (Rosemont, IL) and Novus Biologicals (Burlington, Vt), respectively. Mouse anti-beta actin was from ThermoFisher (Waltham, MA).

#### Time-course study of BDD and N6 expression after DNA vector delivery

Male *C57BL/6J* mice (6-8 weeks old) received pMT, pMT-BDD, or pMT-N6 vector DNA through hydrodynamic tail vein injection described above. Plasma samples were collected at 23-24 h after DNA vector injection via orbital vein bleeding. Mice were sacrificed either at 24 h or 7 days after vector DNA injection to harvest liver and plasma samples. Sodium citrate (final concentration= 0.4%) was used as an anticoagulant for all blood samples. BDD and N6 levels in the plasma samples and liver homogenates were determined using a human FVIII-specific ELISA kit (Affinity Biologicals Inc.). Hepatic levels of BDD and N6 transcripts were measured by quantitative RT-PCR (RT-qPCR) as described^7^ using the following PCR primer pairs for mouse beta-actin (*Actb*) and human F8 (F8), respectively: *Actb*-fw 5’-GGCTGTATTCCCCTCCATCG-3’, *Actb*-rev 5’-CCAGTTGGTAACAATGCCATGT-3’; F8-fw 5’-GCATTCGCAGCACTCTTCG-3’, F8-rev 5’-GAGGTGAAGTCGAGCTTTTGAA-3’. The RNA samples were subjected to DNAse-treatment using a Turbo DNA-free kit (Fisher Scientific, Waltham, MA) to eliminate contamination with pMT-BDD and pMT-N6 vectors before being used for RT-qPCR analysis.

#### Histology and immunohistochemistry staining

Formalin-fixed paraffin-embedded (FFPE) blocks of mouse livers were prepared from empty vector, BDD, and N6-transduced mice. Tissue fixation and paraffin embedding was performed following routine methods. Four μm-thick serial sections were cut from each FFPE block and adhered to charged glass slides (Superfrost Plus; Fisher Scientific, Waltham, MA, USA). Sections were incubated at 60°C for 1 hr prior to deparaffinization in xylene and then rehydrated in 100%, 95%, and double-distilled water successively. Sections were heat-induced in retrieval buffer at pH 6.0, incubated with peroxidase block (Dako, Mississauga, ON, Canada) for 20 min followed by blocking (5% goat serum in 1% PBS-T) for 1 hr. Sections were then incubated overnight at 4°C with primary antibodies diluted in blocking buffer. Primary antibodies used for this study were: rabbit anti-GS (Glutamine Synthetase: Abcam, Cambridge, UK, ab176562; dilution 1:1000), rabbit anti-CD34 (Abcam, Cambridge, UK, ab81289; dilution 1:2500), rabbit anti-CTNNB1 (β-catenin: Abcam, Cambridge, UK, ab223075; dilution 1:4800), rabbit anti-CD44 (Abcam, Cambridge, UK, ab157107; dilution 1:2500), rabbit anti-GPC3 (Glypican-3: LSBio, Seattle, USA, LS-B13373; dilution 1:4000), rabbit anti-BiP (Binding Immunoglobulin Protein: Cell Signaling Technology, Massachusetts, USA, #3177; dilution 1:600). The detection system used was the EnVision+ System-HRP kit (Dako, K4007). Sections were counterstained with hematoxylin prior to dehydration and mounted with Permount (Fisher, SP-15-100). The first section of each series was stained with Hematoxylin and Eosin (H&E) and the fifth section was stained with reticulin (Abcam, #ab150684) for an initial histopathological assessment. Immunohistochemistry with Thio-S and FVIII antibodies was performed as described.^8^

#### Scoring analysis

Slides were scanned using the Aperio AT Turbo system at 20x magnification and viewed using the Aperio Image Scope system. The positivity [Total number of positive pixels divided by total number of pixels: (NTotal – Nn)/(NTotal)] was assessed using the positive pixel count algorithm from the ImageScope software (positive pixel count V9, Aperio, Inc.). The algorithm outputs a strong, moderate, and weak intensity range. The total signal positivity was used to quantify all immunohistochemistry markers where only the strong signal positivity was used as a measure of total positivity of apoptotic cells for TUNEL assay positivity quantification. Signal positivity was analyzed at the central tumor, the interface and distal normal liver.

#### Histopathological analysis

All slides were analyzed, in a blinded fashion, by a board-certified pathologist who evaluated the presence of hepatocellular adenomas and carcinomas based on an immunohistology analysis and various morphometrical features unique to each type.

The following are the stains used to first confirm the lesions and to evaluate whether they represent adenomas (benign disease) or carcinomas. Reticulin staining is intended to demonstrate reticular fibers surrounding tumor cells. Tumors that retain a reticulin network are generally benign or pre-neoplastic, whereas HCC loses the reticulin fiber normal architecture.^9,10^ CD34 is a marker of “capillarization”, a process by which sinusoid endothelial cells lose their fenestration. CD34 demonstrates neovascularization in HCC, while nontumorous hepatic sinusoids do not stain with CD34.^10–12^ However, it was previously reported that in some cases, CD34 may occur diffusely or in areas of hepatocellular adenoma.^9^ Glypican 3 (GPC3) was identified as a tumor marker for the diagnosis of HCC due to its specificity and sensitivity.^10,13–15^ GPC3 shows a cytoplasmic, membranous and canalicular staining among HCCs. However, among livers with a cirrhotic background, GPC3 expression can also be found focally. Although GPC3 is a specific immunomarker for HCC, and show negative staining in adenomas, the diagnosis of HCC should not only rely on this marker.^16,17^ Glutamine synthetase (GS) stains positively for zone 3 of the liver parenchyma. GS overexpression is related to a β-catenin mutation which is a well-known indicator for malignant transformation.^18^ Most HCCs show strong positive cytoplasmic staining while adenomas are known to stain for GS at variable frequencies^14,19,20^. Consequently, GS is a less valuable immunomarker for differentiating HCC from adenomas and alternatively preferred to differentiate between HCC and dysplastic nodules. Beta-catenin shows membranous positive staining on normal hepatocytes. Activation of β-catenin leads to cytoplasmic and nuclear accumulation of β-catenin, which is suggestive of malignant transformation.^21,22^ Finally, CD44 is an adhesion molecule found on sinusoidal lymphocytes and Kupffer cell types in normal liver.^23–25^ Expression of CD44 in HCC cells has been associated with invasive properties.^26^

Apart from the above immunohistology analysis, the lesions categorization into hepatocellular adenoma or hepatocellular carcinoma was further supported based on the identification of 4 morphometrical features. Hepatocellular adenoma was defined as 1. hepatocellular plates 1-2 cells thick 2. nuclear density (#nuclei/mm^2^) <1.5x of adjacent non-neoplastic liver, 3. no obvious nuclear atypia, 4. no pseudogland formation. Hepatocellular carcinoma defined as 1. hepatocellular plates >3 cells thick, 2. nuclear density > or =1.5X of adjacent non-neoplastic liver, 3. obvious nuclear atypia, 4. presence of occasional pseudoglands.^27^

#### Microdissection and DNA analysis of the tumor and non-tumor tissues

Manual microdissection was used to isolate specific regions from FFPE BDD and N6 normal liver and tumor tissue samples. Sectioning was performed by cutting 7 μm thick sections of the FFPE tissue samples using the microtome and sections are mounted on charged glass slides (Superfrost Plus; Fisher Scientific, Waltham, MA, USA). Regions of interest were selected by visually using an inverted light microscope and tissues were manually scraped into 1.5-ml microcentrifuge tubes. Between 6 to 10 tissue sections were collected into each tube and stored at −20°C prior to DNA isolation. FFPE liver blocks prepared from saline- and pMT-BDD-injected mice that were sacrificed at 24 h after hydrodynamic tail vein injection were used as negative and positive controls, respectively, for this experiment. DNA were isolated from the dissected tissues using a QIAamp DNA FFPE Tissue Kit (Qiagen Inc., Valencia, CA) according to the manufacturer’s instruction. Approximately ~0.2 μg DNA from each sample was used for direct qPCR analysis to detect BDD and N6 coding sequences in the tumor and non-tumor tissues using a CFX384 Real-Time System (Bio-Rad, Hercules, CA) described for RT-qPCR.^7^ The same primer pair that was used for RT-qPCR analysis of BDD and N6 transcripts described above (i.e, F8-fw 5’-GCATTCGCAGCACTCTTCG-3’, F8-fw 5’-GAGGTGAAGTCGAGCTTTTGAA-3’) was used for this purpose whereas the following primer pair was used for amplification of mouse beta actin (*Actb*) genomic sequence: *Actb*-genomic-fw: 5’-AACCGTGAAA AGATGACCCAG-3’, *Actb*-genomic-rev: 5’-CACAGCCTGGAT GGCTACGTA-3’.

#### Statistical Analyses

Statistical analyses were performed using GraphPad Prism 9. Chi-square test was used for comparing of tumor incidence between BDD and N6 groups. A p value < 0.05 was regarded as statistically significant. Statistical analysis for immunohistochemistry was performed with a two-tailed Student’s *t*-test. *P* values of <0.05 were considered significant (p < 0.05 *, p < 0.01 **, p < 0.001 ***).

## Supplemental Data

**Figure S1.**
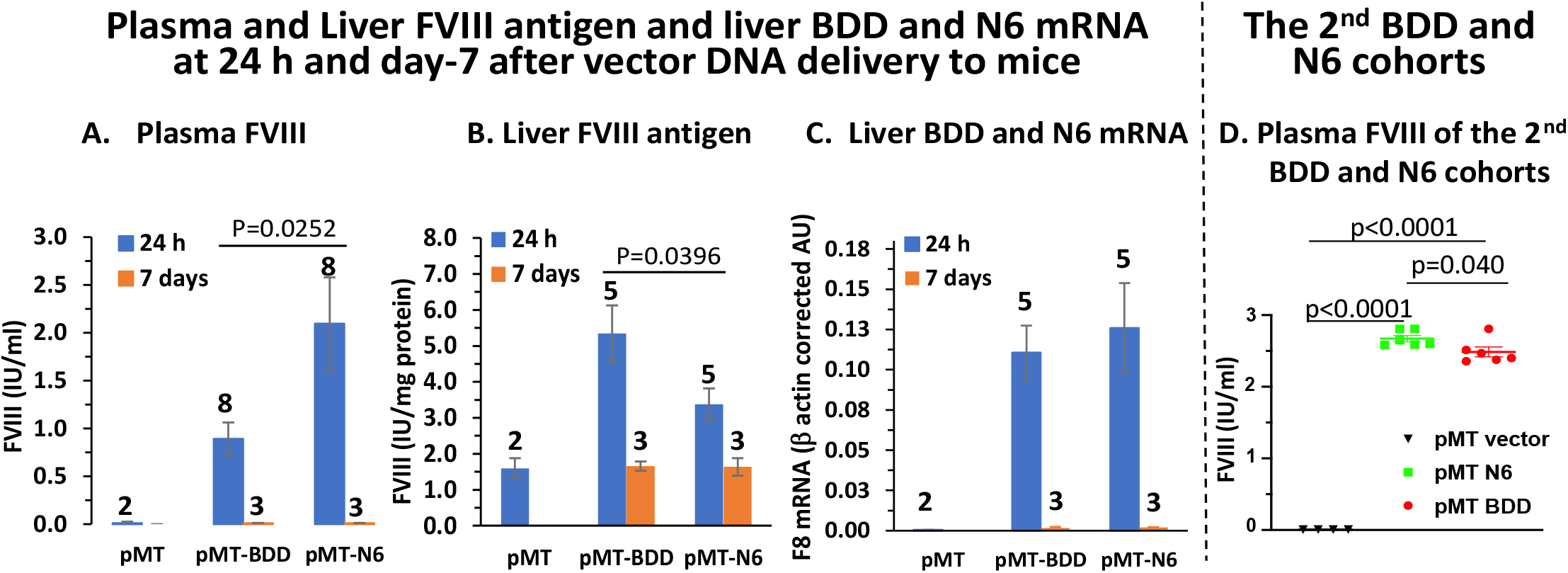
Plasma and Liver FVIII antigen and hepatic BDD and N6 transcripts were undetectable 7 days after vector DNA delivery to mice. **A)** Plasma BDD and N6 antigen. **B)** Liver human FVIII antigen. **C)** Liver BDD and N6 mRNA levels. Mice were sacrificed at 24 h or 7 days after hydrodynamic tail injection of pMT, pMT-BDD or pMT-N6 DNA vectors. Data are represented as mean ± S.E.M. The numbers on the top of each bar in the graphs of panels A-C are the number of mice (n) for the corresponding data. **D)** Plasma antigen levels of BDD and N6 for the second cohort of TM-BDD and TM-N6 mice that were used for the tumor study described in Figures 6 and S2. Plasma samples from these mice were also obtained at 24 h after vector DNA injection. Plasma BDD and N6 antigen levels in these mice appeared to be higher levels than those shown in Figures 6A. This difference might have resulted from variable nature of the DNA delivery, ELISAs performed at different times, as well as murine housing conditions.

**Figure S2.**
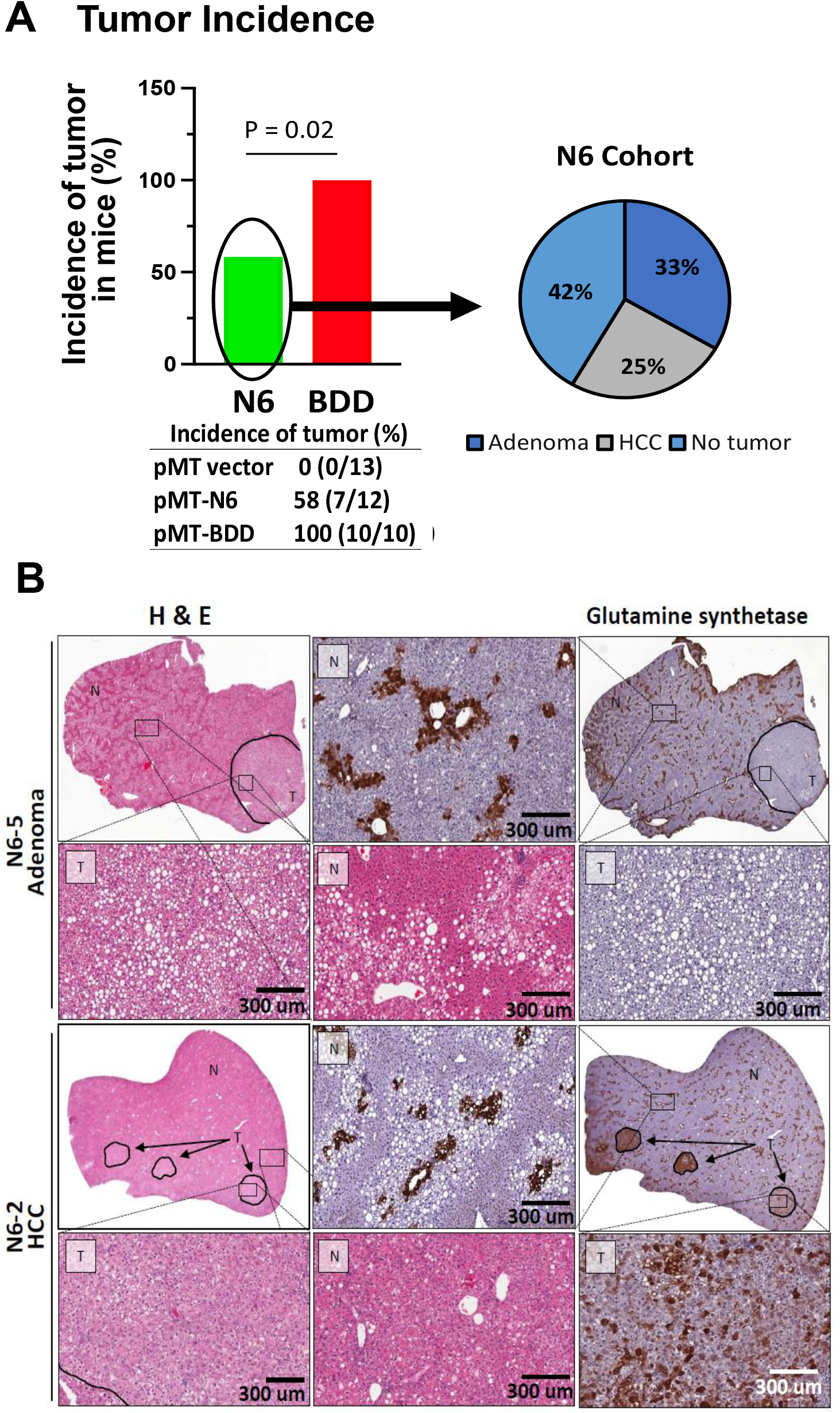
Liver surface tumors in mice injected with N6 vector DNA. Mice were injected with BDD, N6, or empty vector DNA and treated with a high-fat diet as described in Methods and legend to Fig. 6. **A)** Distribution of HCC and adenomas among liver tumors in the N6 cohort. **B)** Positivity for glutamine synthase stains was observed in HCC but not in adenoma of the N6 mice. T=tumor, NT=non-tumor.

**Figure S3.**
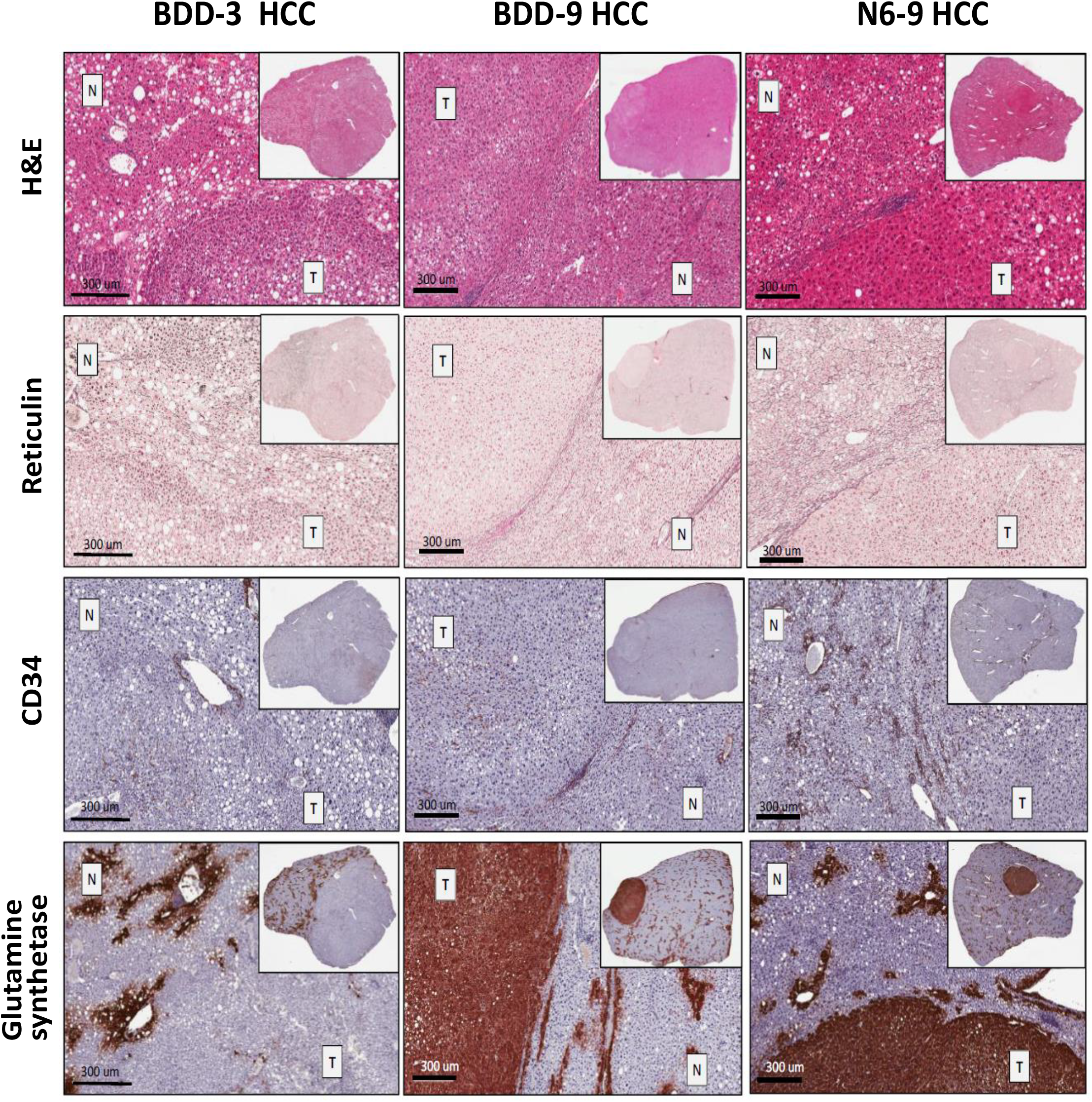
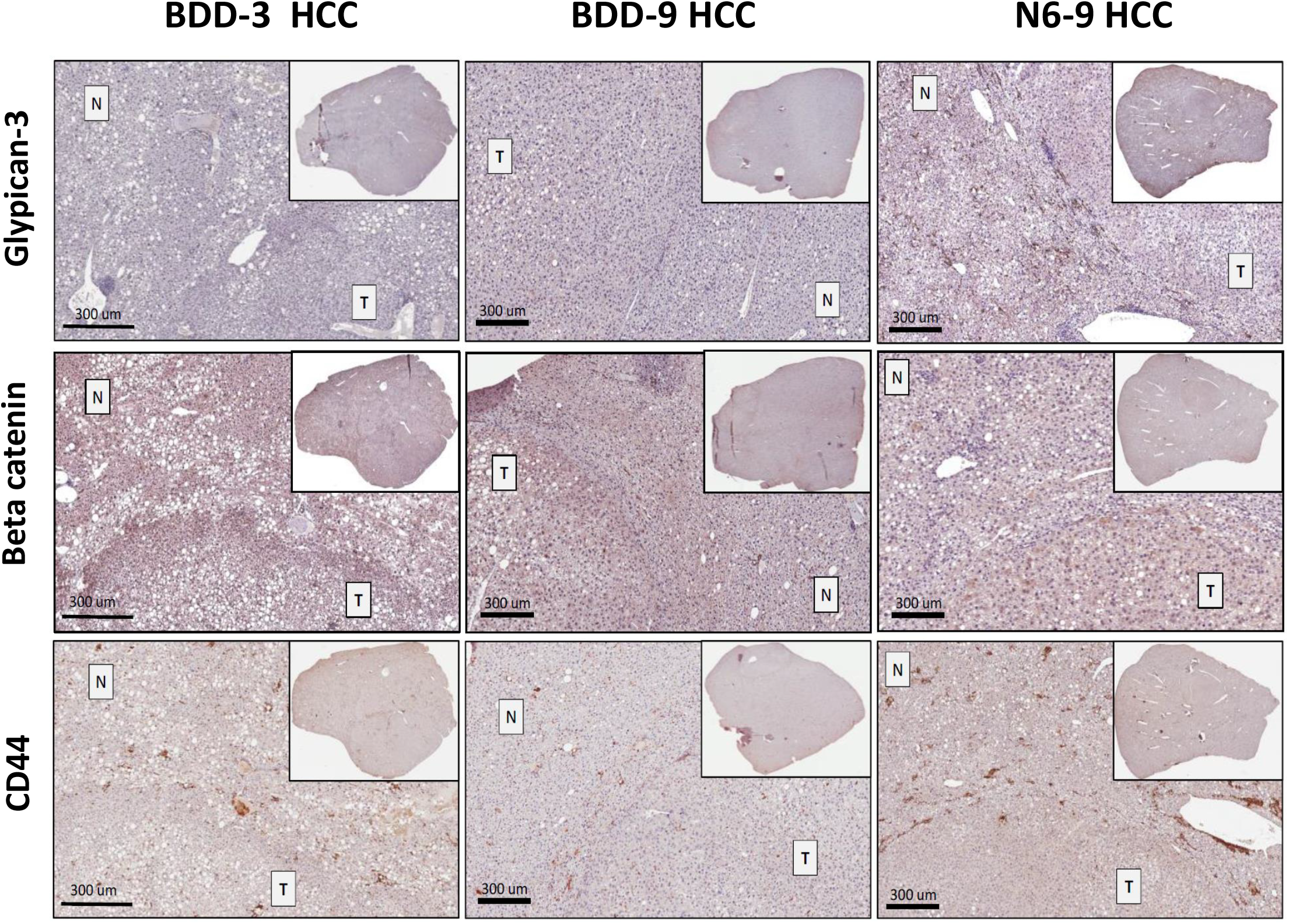
Representative immunohistochemistry characterization of HCC samples. Serial sections were stained for H&E: Hematoxylin and Eosin, reticulin, CD34, glutamine synthetase, glypican-3, β-catenin, and CD44. T=tumor, NT=non-tumor. Serial sections were stained for glypican-3, β-catenin, and CD44. T=tumor, NT=non-tumor.

**Table S1.**
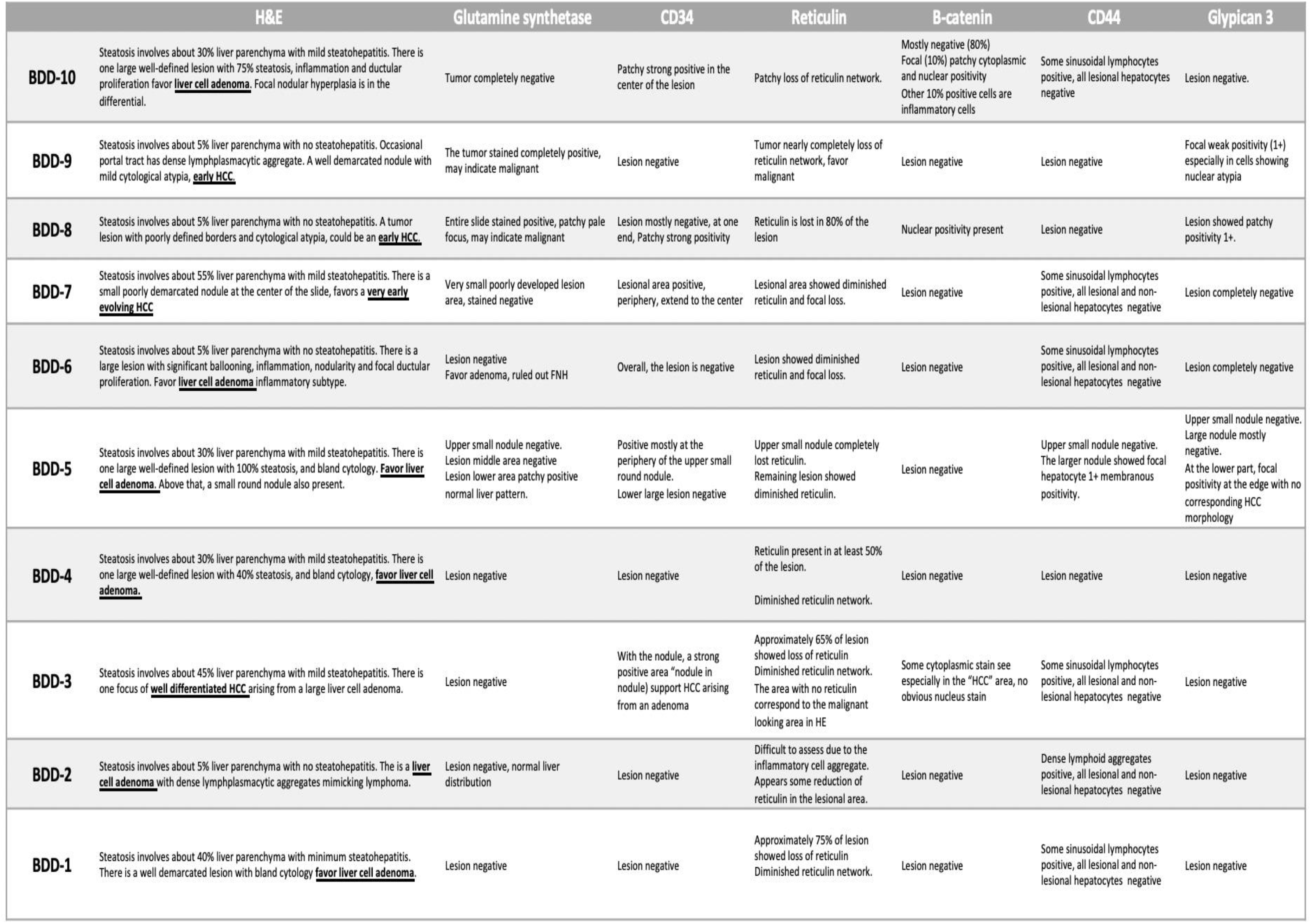
Immunohistochemistry characterization of the pMT-BDD cohort samples.

**Table S2.**
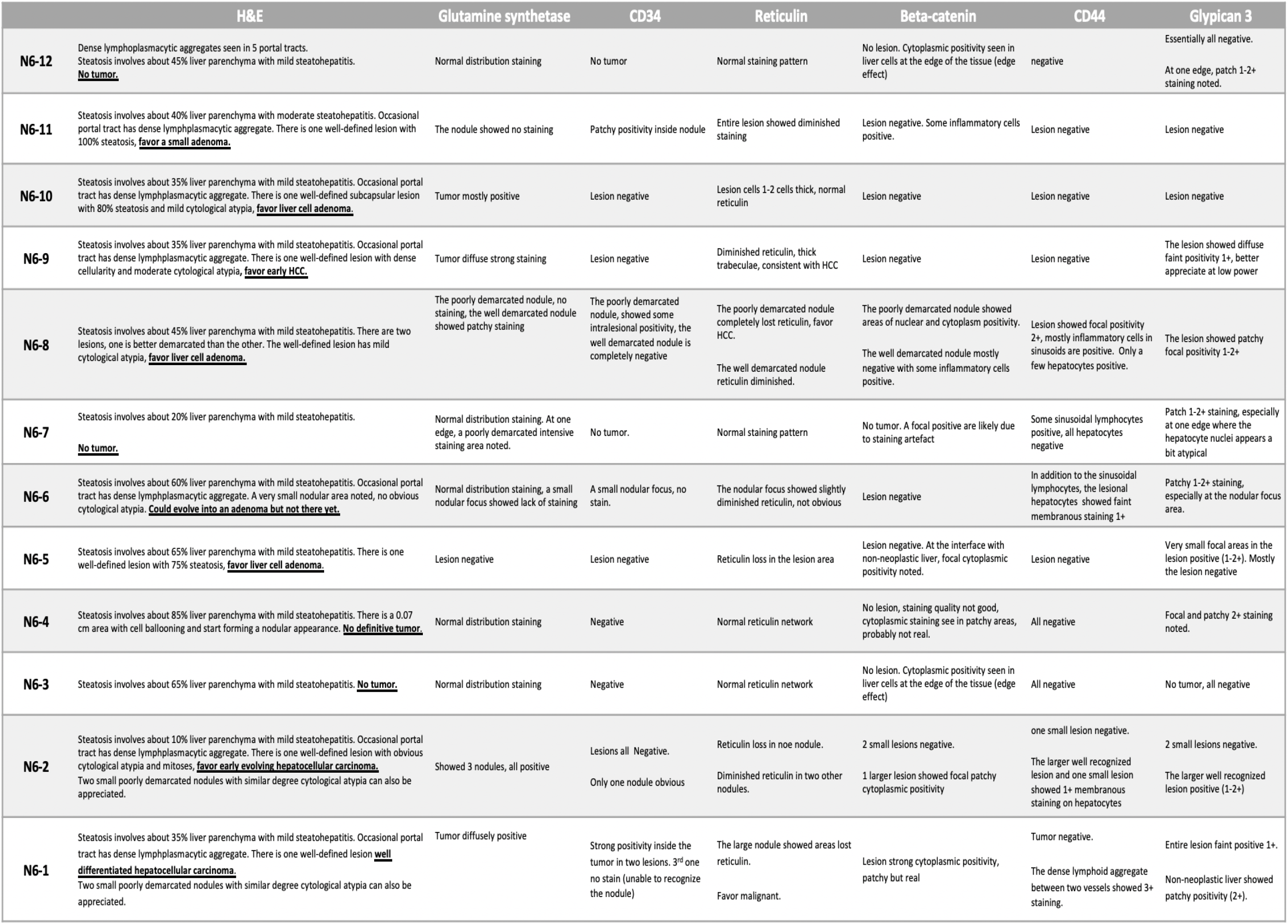
Immunohistochemistry characterization of the pMT-N6 cohort samples.

**Table S3.** Immunohistochemistry scores of tumor markers in the BDD and N6 cohort samples. **Scoring Interpretation-*Reticulin*:** Reticulin lost is suggestive of malignancy.^9,10^ The progressive lost of associated reticulin fiber is labeled as follows: 3+: Intact reticulin pattern (1-2 cells thickening of hepatic plates), 2+: Weak (2-3 cells thickening of hepatic plates), 1+: Moderate (>4 cells thickening of hepatic plates), and Neg: Complete loss of the reticulin. ***Glypican-3*:** Cytoplasmic granular staining supports malignancy.^10,14,15^ Presence of cytoplasmic diffused or patchy and/or focal patterns are labeled with different intensity as follows: 3+: Strong positivity, 2+: Moderate positivity, 1+: Weak positivity, and Neg: No Positive staining. ***CD34*:** Diffused and strong sinusoidal endothelial cell staining is observed in almost all HCC cases.^10,12^ The endothelial cells of the sinusoidal wall are labeled as follows: 2+: Diffusely Positive (>80% endothelial cells of the sinusoidal wall), 1+: Focally Positive (30-80% endothelial cells of the sinusoidal wall), and Neg: Negative (<30% endothelial cells of the sinusoidal wall). ***Glutamine Synthetase*:** HCCs tumors show strong positive staining while adenomas may occasionally show positive staining.^14,19,20^ Lesions are labeled as follows: Pos. Diffused: Diffused cytoplasmic positivity, Pos. Patchy: Patch like cytoplasmic positivity, Pos. Weak: Weak cytoplasmic positivity, and Neg: No positive staining. In normal liver, glutamine synthetase is confined in 2-3 cell-thick hepatocytes around the central vein and is labeled as: Pos. CV. ***β-Catenin*:** Nuclear accumulation of β-catenin is suggestive of malignant transformation.^21,22^ Presence of cytoplasmic and/or nuclear staining in cancer cells are labeled as follows: Diffused: Diffused cytoplasmic Positivity, Nuclear: Nuclear positive staining, Diffused & Nuclear: Diffused cytoplasmic and nuclear positivity, and Neg: No Positive staining. ***CD44*:** Upregulation of CD44 expression is associated with HCC development.^28,29^ Diffused or focal staining patterns of CD44 in cancer cells or hepatocytes is labeled as follows: 3+: Strong membranous positive cells, 2+: Moderate membranous positive cells, 1+: Weak membranous positive cells, and Neg: No positive staining.

| Liver Samples | # of Tumors | Lesions Type | Reticulin | Glypican-3 | CD34 | Glutamine Synthetase | β-catenin | CD44 |
| --- | --- | --- | --- | --- | --- | --- | --- | --- |
| BBD-1 | 1 | Adenoma | 2+ | Neg | Neg | Neg | Neg | Neg |
| BBD-2 | 1 | Adenoma | 3+ | Neg | Neg | Neg | Neg | Neg |
| BBD-3 | 1 | HCC | Neg | Neg | 2+ | Neg | Cytoplasmic | Neg |
| BBD-4 | 1 | Adenoma | 2+ | Neg | Neg | Neg | Neg | Neg |
| BBD-5 | 1 | Adenoma | 2+ | Neg | Neg | Pos, Patchy | Neg | Focal, 1+ |
| BBD-6 | 1 | Adenoma | 3+ | Neg | Neg | Neg | Neg | Neg |
| BBD-7 | 1 | HCC | 1+ | Neg | 1+ | Neg | Neg | Neg |
| BBD-8 | 1 | HCC | 1+ | Patchy, 1+ | 2+ | Pos, Diffused | Nuclear | Neg |
| BBD-9 | 1 | HCC | Neg | Focal, 1+ | Neg | Pos, Diffused | Neg | Neg |
| BBD-10 | 1 | Adenoma | 2+ | Neg | 1+ | Neg | Cytoplasmic & Nuclear | Neg |
| N6-1 | 3 | HCC | Neg | Diffused, 2+ | 2+ | Pos, Diffused | Cytoplasmic | Neg |
| N6-2 | 3 | HCC | 2+ | Diffused, 1-2+ | 2+ | Pos, Diffused | Cytoplasmic | Focal, 1+ |
| N6-3 | 0 | N/A | 3+ | Neg | Neg | Pos, CV | Neg | Neg |
| N6-4 | 0 | N/A | 3+ | Patchy & Focal, 2+ | Neg | Pos, CV | Neg | Neg |
| N6-5 | 1 | Adenoma | 2+ | Neg | Neg | Neg | Cytoplasmic | Neg |
| N6-6 | 0 | N/A | 3+ | Patchy, 1-2+ | Neg | Pos, CV | Neg | Weak, 1+ |
| N6-7 | 0 | N/A | 3+ | Patchy, 1-2+ | Neg | Pos, CV | Neg | Neg |
| N6-8 | 2 | Adenoma | 2+ | Patchy & Focal, 1-2+ | Neg | Pos, Patchy | Neg | Focal, 2+ |
| N6-9 | 1 | HCC | Neg | Diffused, 1+ | Neg | Pos, Diffused | Neg | Neg |
| N6-10 | 1 | Adenoma | 3+ | Neg | Neg | Pos, Diffused | Neg | Neg |
| N6-11 | 1 | Adenoma | 2+ | Neg | 1+ | Neg | Neg | Neg |
| N6-12 | 0 | N/A | 3+ | Neg | Neg | Pos, CV | Neg | Neg |

**Table S4.** BDD and N6 DNA in tumor and non-tumor tissues in livers of cohorts of pMT-BDD and pMT-N6 injected mice was not detectable by quantitative PCR analysis. DNAs were isolated from the tumor and non-tumor tissues of FFPE liver blocks of mice listed below and subjected to qPCR analysis as described in Methods, with mouse beta actin (*Actb*) as an internal control samples to normalize the data for BDD and N6 vector sequence detection [F8/*Actb*= 2^(Cq-*Actb* – Cq-F8)]. Except for the positive control DNA (*i,e*., pMT-BDD at 24h) sample (F8/*Actb* = 11.17), the F8/*Actb* values for all other samples were within the range of background noise, indicating the absence of the BDD or N6 sequence. NA: non-applicable; ND: not determined due to insufficient amounts of tissues; NV: no value /below detection limit.

|  |  | Quantification cycle (Cq) |  |  |  |  |  |
| --- | --- | --- | --- | --- | --- | --- | --- |
|  |  | Tumor |  |  | Non-tumor |  |  |
| | Sample ID | F8 | *Actb* | $F8/Actb$ | F8 | *Actb* | $F8/Actb$ |
| Negative control | Saline-24h | NA | NA | NA | 27.004 | 21.427 | 0.021 |
| Positive control | pMT-BDD-24h | NA | NA | NA | 17.771 | 21.253 | 11.169 |
| pMT-BDD cohort | BDD-1 | 35.745 | 27.175 | 0.003 | 33.470 | 26.950 | 0.011 |
|  | BDD-2 | NV | 30.958 | NV | 34.244 | 31.244 | 0.125 |
|  | BDD-3 | NV | 25.997 | NV | NV | 26.540 | NV |
|  | BDD-4 | NV | 30.634 | NV | 38.436 | 31.159 | 0.006 |
|  | BDD-5 | 37.637 | 27.746 | 0.001 | NV | 28.466 | NV |
|  | BDD-6 | 33.283 | 27.094 | 0.014 | 35.302 | 25.668 | 0.001 |
|  | BDD-7 | ND | ND | ND | 34.860 | 26.278 | 0.003 |
|  | BDD-8 | 33.836 | 26.908 | 0.008 | ND | ND | ND |
|  | BDD-9 | 34.006 | 26.091 | 0.004 | 35.160 | 27.183 | 0.004 |
|  | BDD-10 | 33.373 | 29.238 | 0.057 | 34.749 | 30.486 | 0.052 |
| pMT-N6 cohort | N6-1 | 35.274 | 28.220 | 0.008 | 35.665 | 26.535 | 0.002 |
|  | N6-2 | 33.325 | 29.441 | 0.068 | NV | 32.852 | NV |
|  | N6-3 | NA | NA | NA | 34.267 | 29.593 | 0.039 |
|  | N6-4 | NA | NA | NA | 36.750 | 26.847 | 0.001 |
|  | N6-5 | 38.596 | 34.724 | 0.068 | NV | 28.666 | NV |
|  | N6-6 | NA | NA | NA | NV | 29.190 | NV |
|  | N6-7 | NA | NA | NA | 34.546 | 27.072 | 0.006 |
|  | N6-8 | 37.233 | 29.000 | 0.003 | 36.919 | 27.365 | 0.001 |
|  | N6-9 | 36.809 | 28.160 | 0.002 | 36.084 | 30.111 | 0.016 |
|  | N6-10 | 34.615 | 26.158 | 0.003 | 34.491 | 26.627 | 0.004 |
|  | N6-11 | NV | NV | NV | 34.421 | 26.518 | 0.004 |
|  | N6-12 | NA | NA | NA | 33.894 | 29.693 | 0.054 |

